# Population dynamics of *Pelagibacterales* clonal lineages: ecological-consortia and frequency modulation

**DOI:** 10.64898/2026.06.30.735289

**Authors:** Ana-Belen Martin-Cuadrado, Jose M. Haro-Moreno, Francisco Rodriguez-Valera

**Affiliations:** Department of Physiology, Genetics and Microbiology, Universidad de Alicante, Alicante, Spain; Evolutionary Genomics Group, División de Microbiología, Universidad Miguel Hernández, Alicante, Spain

**Keywords:** SAR11, Pelagibacterales, clonal diversity, pangenome, Mediterranean Sea, metagenomics, log-normal abundance

## Abstract

*Pelagibacterales* gMED is the dominant epipelagic genomospecies in the western Mediterranean Sea. We used the O-chain biosynthesis gene clusters, OBCs, as clonal barcodes to analyse strain-level population structure. In total, 385 OBC-defined clonal lineages were tracked across Mediterranean metagenomes spanning 14 years and depths from 5 to 90 m within the photic zone, with between 128 and 336 detected per metagenome. The relative conservation of dominant OBC types across years, seasons, and geographic locations indicated a persistently high and stable clonal diversity. Abundance distributions were log-normal in all metagenomes — the canonical signature of bet-hedging portfolio dynamics. Co-varying consortia of clonal lineages were detected, identifying ecologically coherent guilds whose relative abundances fluctuate in concert across environmental gradients, particularly depth during the stratified season. Consortia membership showed no correlation with core-genome phylogeny, indicating convergent ecological adaptation independently of ancestry. Our work reveals another layer of complexity below the community level, which together with its modulation, provides the gMED population with a robust and versatile genetic repertoire for the degradation of dissolved organic matter under the fluctuating conditions of the epipelagic ocean. We have named this mechanistic model frequency-modulation (FM).

## Introduction

Different approaches have established that in natural aquatic habitats, the population of a bacterial species is composed of multiple clonal lineages, each characterised by a distinct flexible gene pool (Gonzaga et al. 2012, Kashtan et al. 2014, Viver et al. 2024, Haro-Moreno et al. 2025, Molina-Pardines et al. 2025). Two complementary mechanisms have been proposed to account for the maintenance of high clonal diversity in free-living aquatic bacteria (Rodriguez-Valera and Molina-Pardines 2026). First, the ‘constant-diversity’ (CD) hypothesis (Rodriguez-Valera et al. 2009) posits that phages keep all clones in dynamic equilibrium through negative frequency-dependent selection: as any clone increases in frequency, its cognate phages proliferate and cull it, allowing rarer clones to recover. Second, clones may partition resources — for example, by exploiting the DOM released by phage lysis of fast-growing clones (Wilhelm 1999), or by exchanging nutrients through extracellular vesicles (Biller et al. 2014, Biller et al. 2022) — thereby decoupling fitness from the growth rate of individual cells (Rodriguez-Valera and Molina-Pardines 2026).

If such clonal consortia are a general feature of free-living marine bacteria, the high diversity found at any given time and location would be expected to remain relatively stable, despite fluctuating conditions. The relative abundance of individual clones may nevertheless respond to environmental gradients, particularly the vertical (depth) stratification and, to a lesser extent, horizontal heterogeneity of the water masses (Li et al. 2026). For photoheterotrophs such as most Pelagibacterales, these physical structures are critical determinants of ecological fitness. Here we analyse the proportions of different clonal lineages across time and depth in the gMED genomospecies (*Pelagibacterales* subclade Ia.3/VII; (Haro-Moreno et al. 2020), now assigned to the genus *Angustipelagibacter* (Freel et al. 2025), which dominates epipelagic recruitment in Mediterranean Illumina metagenomes (Haro-Moreno et al. 2020). We leveraged a large collection of long-read (PacBio) metagenome-derived fragments of the O-chain biosynthesis gene cluster as a high-resolution clonal barcode. This hypervariable surface-polysaccharide *locus* has been exploited as a strain marker in Gram-negative pathogens for over a century, most notably in the Enterobacteriales (Holt et al. 2020), and is known to encode major phage-receptor determinants among other phenotypic markers (May et al. 2025). Its utility for characterising clonal lineages that vary in their flexible genome (flexome) content, across the *Pelagibacterales* clade has been established (Haro-Moreno et al. 2025). Long-read metagenomes provide partial OBC sequences from uncultured cells that, although often incomplete, allow identification of the different clonal lineages (strains). We have also analyzed single-amplified genomes (SAGs) We also analysed single-amplified genomes (SAGs) that, although scarcer, provide more complete reference OBCs. Coverage and read recruitment at high nucleotide identity (≥97%) of Illumina metagenomes against these OBC sequences were used as proxies for the relative prevalence of each clonal lineage. Overall, our results support the relative stability of the clonal structure of gMED populations, favouring ecological equivalence and resource sharing over periodic selective sweeps that would decrease diversity or dramatically alter population structure (Bendall et al. 2016). Furthermore, the detection of co-varying clone consortia among them—groups that fluctuate in a coordinated fashion across environmental gradients— points to a novel unit of intraspecies diversity, distinct from subspecies or ecotypes (Kujawinski et al. 2023) and more akin to the pathotypes of pathogenic bacteria (Geurtsen et al. 2022) in which flexible genes determine infectivity.

## Results

We collected 354 partial O-chain biosynthesis gene clusters from six long-read metagenomes (LR-OBCs) retrieved from a station 20 NM off the coast of Alicante, Spain (Supplementary Table 1). These OBCs were assigned to the gMED genomospecies based on the 16S–23S internal transcribed spacer (ITS), which in this species is consistently adjacent to the OBC locus (Giovannoni et al. 2005, Tripp et al. 2011, Haro-Moreno et al. 2020, Haro-Moreno et al. 2025). We also included in the analysis a set of 31 SAGs from the Bermuda Atlantic Time-series/Sargasso Sea collection (SAG-AG) (Pachiadaki et al. 2019) and from a single north-western Mediterranean sample (SAG-MED) (Haro-Moreno et al. 2020) all within the same gMED genomospecies as described previously (Haro-Moreno et al, 2020). The 354 LR-OBCs were recovered from PacBio metagenomes retrieved at the same Mediterranean off-Alicante site and therefore represent abundant local lineages in the South-Western Mediterranean. The 31 SAG-OBCs, by contrast, were drawn from geographically diverse collections and thus could represent a broader cross-section of global gMED diversity, including lineages that might be rare or absent at the metagenomes analyzed.

### Relative clone abundances per metagenome

A total of 27 short-read (Illumina) metagenomes from the Mediterranean were used in this analysis, ten of them from the TARA Oceans collection, spanning 2009-2023, different locations, and depths from 5 to 90 m within the photic zone (Supplementary Table 1). Read recruitment normalised by reference sequence length and metagenome size (reads per Kb of genome and Gb of metagenome, RPKG) was used as the primary abundance metric throughout (Supplementary Table 2; see Methods). On average of 264.3 ± 54.2 different LR-OBCs were detected per near-surface metagenome across the seven off-Alicante 15–20 m samples (range 157-327; 74.7 ± 15.30% of the 354 LR-OBCs). The number of Illumina fragments recruited at ≥97% nucleotide identity per OBC varied from 3 to 11,314 across all 27 metagenomes (median 410; mean 734 ± 1,001). Detection increased from 44.4% in September 2014 (15 m) to 92.4% in the Winter-2022 (20 m) mixed-water sample. Strikingly, only 2 LR-OBCs were absent from all seven near-surface samples, while 118 OBCs (30.6% of the full OBC set, including 100 LR-OBCs and 18 SAG-OBCs) were detected in every one of them, and 328 (85.2%) appeared in four or more (Fig. 1). The detection rate of all OBC–metagenome combinations was 68%, remarkably high considering the time and space gap separating the metagenomes.

**Fig. 1:**
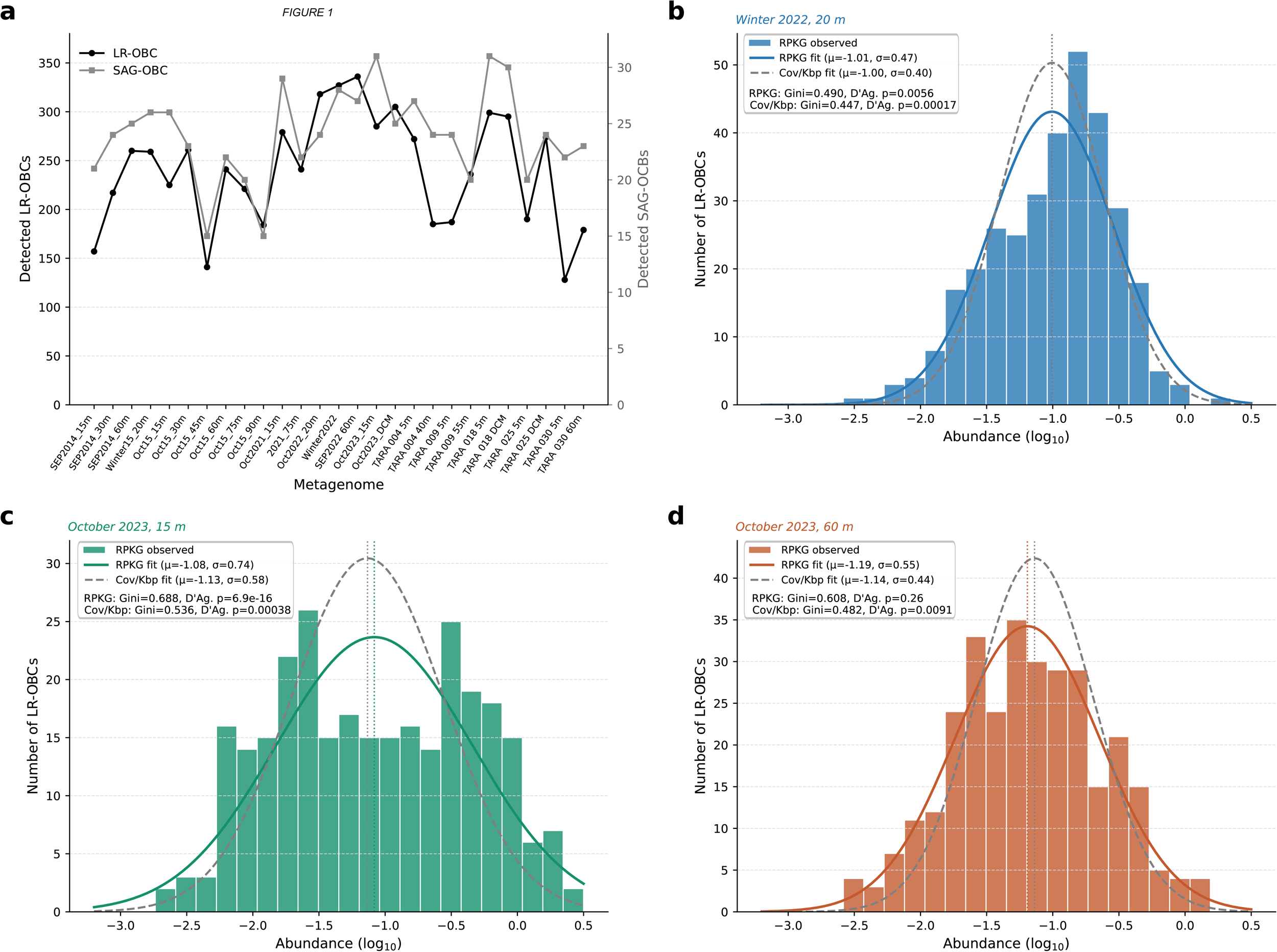
Detection and abundance structure of gMED OBC-defined clonal lineages across Mediterranean metagenomes. **a,** Number of detected OBC-defined clonal lineages per metagenome using vertical RPKG recruitment, shown separately for LR-OBCs and SAG-OBCs. LR-OBCs correspond to O-chain biosynthesis cluster fragments recovered from PacBio HiFi long reads, whereas SAG-OBCs correspond to OBCs retrieved from single-amplified genomes. Detection was defined as non-zero recruitment. **b–d,** Log₁₀-transformed RPKG abundance distributions of LR-OBCs in three representative metagenomes: winter mixed water column at 20 m (b), stratified upper photic layer at 15 m (c), and stratified DCM at 60 m (d). Bars indicate the number of LR-OBCs within each log₁₀(RPKG) bin. Solid black lines show fitted normal distributions in log₁₀ space, and dashed vertical lines indicate the mean log₁₀(RPKG). Reported values correspond to the mean and standard deviation of log₁₀-transformed non-zero RPKG values, the geometric mean RPKG, and the D’Agostino–Pearson omnibus normality test.

Individual LR-OBCs showed broad abundance distributions, consistent with an approximately log-normal structure rather than clonal monopoly (Fig. 1b–d; Supplementary Fig. 1). For the Winter-2022-20 m sample —the highest recruiting metagenome, with 327 LR-OBCs detected — log-transformed RPKG values yields a near-normal distribution (μ = −1.01, geometric mean 0.098 RPKG, σ = 0.47). The D’Agostino–Pearson test formally rejected strict log-normality (p = 0.0056), although the fit was visually close and the deviation was mainly driven by the tails rather than the central distribution. Similarly broad, approximately log-normal distributions were recovered across all conditions examined, from winter mixed waters to stratified surface and DCM layers (Supplementary Fig. 1), and in all cases remained far from the monopoly expected under a clonal-sweep scenario. This pattern — many coexisting lineages spanning nearly three orders of magnitude in abundance, but with a large majority at intermediate values — is the canonical signature of bet-hedging at the within-species level (Schindler et al. 2010), and is consistent with a persistent, diverse clonal pool in which rank switching rather than presence–absence turnover drives variability across samples.

### Rank shuffling within a stable and diverse strain pool

We next focused on the near-surface photic zone (15–20 m depths), where gMED is more abundant, analysing seven samples spanning 2014–2023 retrieved from the South-Western Mediterranean (most come from the off-Alicante station; bottom depth 200 m; 20 NM from Alicante) with the only exception of Oct 2015-20m sample, collected from a different location at 200 NM from Alicante and over 2,000 m depth (Haro-Moreno et al. 2018), thereby minimizing depth and geography effects. RPKG-based rank trajectories across these metagenomes are shown in Fig.2a, with complete detection frequencies and recruitment summaries provided in Supplementary Table 3.

Of the 120 LR-OBCs reaching the top 30 in at least one sample, across 630 pairwise transitions originating from a top-30 position, the probability of a ≥10-fold rank decline was only 36.5% (230/630), indicating that approximately only one in three top-30 appearances was followed by a substantial rank collapse in another sample (Fig.2b). Rank declines were strongest among the highest-ranked lineages: LR-OBCs starting at ranks 1–5 declined ≥10-fold in 69.5% of comparisons, those at ranks 6–10 in 53.3%, those at ranks 11–20 in 31.9%, and those at ranks 21–30 in 16.2%. For broader thresholds, the probabilities were 60.8% for ≥5-fold, 15.6% for ≥20-fold, and 5.2% for ≥50-fold rank declines. Overall, these data indicate substantial stability of the clonal community structure.

Per-OBC volatility level was heterogeneous (Fig.2b and c). A subset of LR-OBCs showed no ≥10-fold rank declines despite repeatedly entering the top-30 pool — the most stable highly-ranked lineages are colored in Fig 2a and were all C1-surface consortium members (see below). R809821 (blue line in Fig 2a) was the most recurrent lineage overall, appearing in five of the seven samples with ≥10-fold declines in only 22.2% (4/18) of comparisons. In contrast, R634316 (red line in Fig 2a) reached rank 1 in two samples but declined ≥10-fold in 75.0% (3/4) of comparisons, illustrating transient dominance. At the other extreme, several LR-OBCs declined ≥10-fold in all valid top-30 starts, achieving transient high abundance in specific samples but remaining essentially absent in others, some examples are: R631669 and R754915 (C1); R128014 and R204472 (C2).

**Fig. 2:**
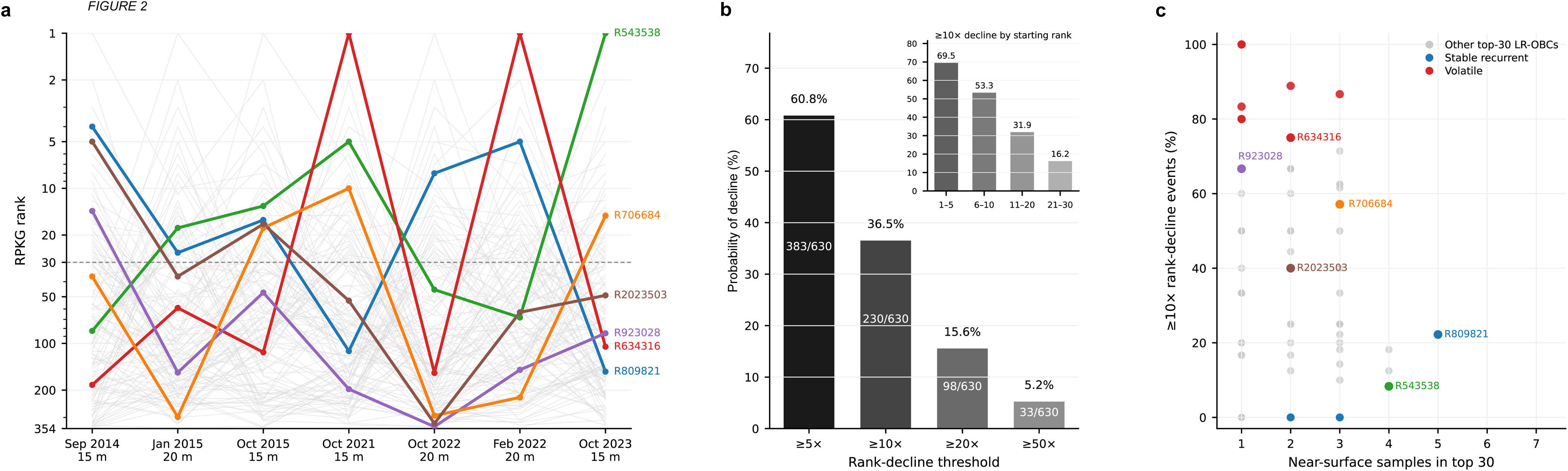
Recurrence and rank reshuffling of high-abundance LR-OBC lineages. **a,** RPKG-based rank trajectories of the 120 LR-OBC lineages that reached the top 30 in at least one of seven near-surface Mediterranean metagenomes sampled between 2014 and 2023. Grey lines indicate all top-30 lineages; coloured lines highlight representative recurrent and transiently dominant lineages. The dashed line marks the top-30 threshold. **b,** Probability of rank decline from a top-30 starting position across 630 valid pairwise comparisons. Bars show ≥5-, ≥10-, ≥20- and ≥50-fold rank declines; numbers indicate transition counts. The inset shows ≥10-fold rank-decline frequency stratified by starting-rank class. **c,** Relationship between recurrence and volatility among top-ranked LR-OBC lineages. Grey points denote all other top-30 lineages; coloured points highlight representative stable recurrent and transiently dominant lineages.

At the per-SAG level, 11 of 31 SAG-OBCs never reached the top 10 in any near-surface sample. Among the 20 that did, seven showed no ≥3-fold decline events. Interestingly, all but one were derived from Atlantic samples (Supplementary Fig. 2). The most volatile were SAG-MED44 (North-Western Mediterranean SAG), with 13 of 17 valid top-10 starts resulting in a ≥3-fold decline (76.5%), and AG-337-J22 (Atlantic), with 12 of 15 such events (80.0%).

Together, both reference sets (LR-OBCs and SAGs) support a model in which a recurrent pool of high-abundance OBC-defined lineages persists through time while dominance is repeatedly redistributed among its members — consistent with rank reshuffling within a stable strain pool rather than sequential clonal replacement or succession.

### Covariation consortia reveal collective frequency modulation

We next asked whether OBC-defined clonal lineages fluctuate independently across metagenomes, or whether subsets of lineages (henceforth called consortia) respond coherently to the same environmental structure. To address this, we removed differences in absolute abundance by applying row-wise Z-score standardisation to log-transformed RPKG values. This transformation retains only the relative recruitment pattern of each lineage across metagenomes, allowing highly and weakly recruited OBCs to be compared according to the shape of their cross-metagenome response.

The LR-OBC-defined strains abundance-independent recruitment profiles resolved into two major covariation consortia (Fig.3a, Supplementary Table 4) (global silhouette coefficient = 0.289; Fig. 3b). Hierarchical clustering using Pearson correlation distance and average linkage identified a surface-associated consortium, C1 (n = 197), characterized by positive Z-scores in surface (5–20 m) metagenomes and negative Z-scores in deep and DCM samples. The second consortium, C2 (n = 157), showed the opposite pattern, displaying preferential recruitment in deep and DCM-associated metagenomes, particularly TARA_009-55 m and MedDCM-SEP2022-60 m, while diminished in surface waters. The two consortia were clearly separated in ordination space (Fig.3c), and k = 2 was the statistically optimal partition across k = 2 to 8 (Fig. 3b). To confirm that this two-consortium structure is not affected by the inclusion of geographically distant TARA Oceans stations, we repeated the clustering analysis using only the 17 Mediterranean metagenomes from the off-Alicante time series (excluding all TARA samples) and the same two-consortium partition was recovered (k = 2, silhouette = 0.372; Supplementary Fig. 3; Supplementary Fig.4; Supplementary Table 5a). These results confirm that the surface/deep consortium structure is a robust feature of the local gMED population and is not driven by the contrast between Mediterranean and Atlantic-influenced TARA stations.

**Fig. 3:**
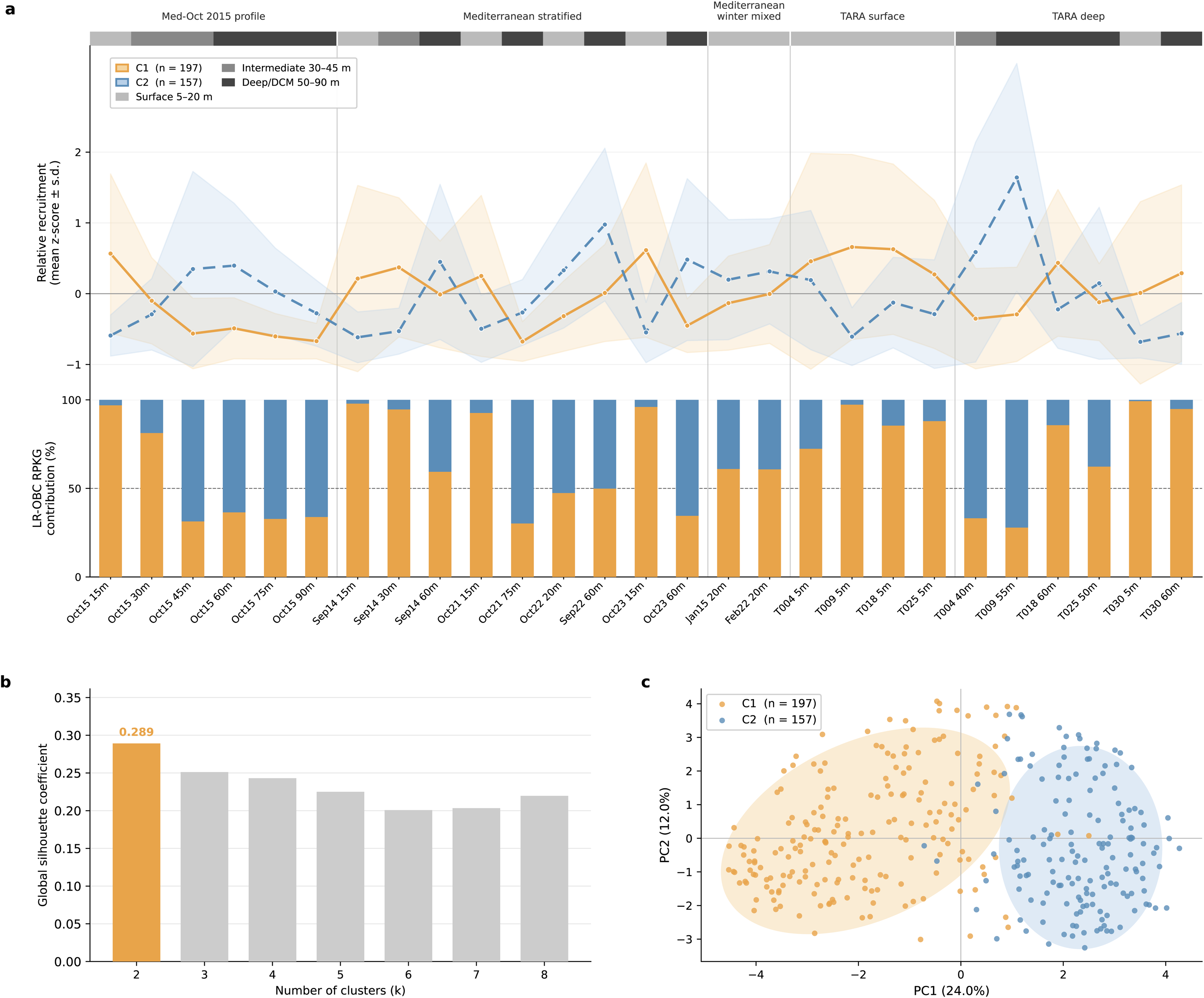
Two major ecological consortia structure LR-OBC recruitment across Mediterranean metagenomes. **a,** *Upper panel*: mean relative recruitment profiles (mean Z-score ± s.d.) of the two major LR-OBC consortia identified by hierarchical clustering of log(1+RPKG)-transformed recruitment values using correlation distance and average linkage. Samples are ordered by hydrographic regime and geographical origin. Shaded areas represent ±1 standard deviation. The coloured bar above the plot indicates depth category (light grey, surface 5–20 m; mid grey, intermediate 30–45 m; dark grey, deep/DCM 50–90 m). *Lower panel*: relative contribution of each consortium to total LR-OBC RPKG per metagenome. **b,** Global silhouette coefficient for clustering solutions k = 2 to k = 8. The optimal partition is highlighted in orange. **c,** Principal component analysis (PCA) of Z-score recruitment profiles. Shaded ellipses represent 1.8 standard deviations around each cluster centroid.

**Fig. 4:**
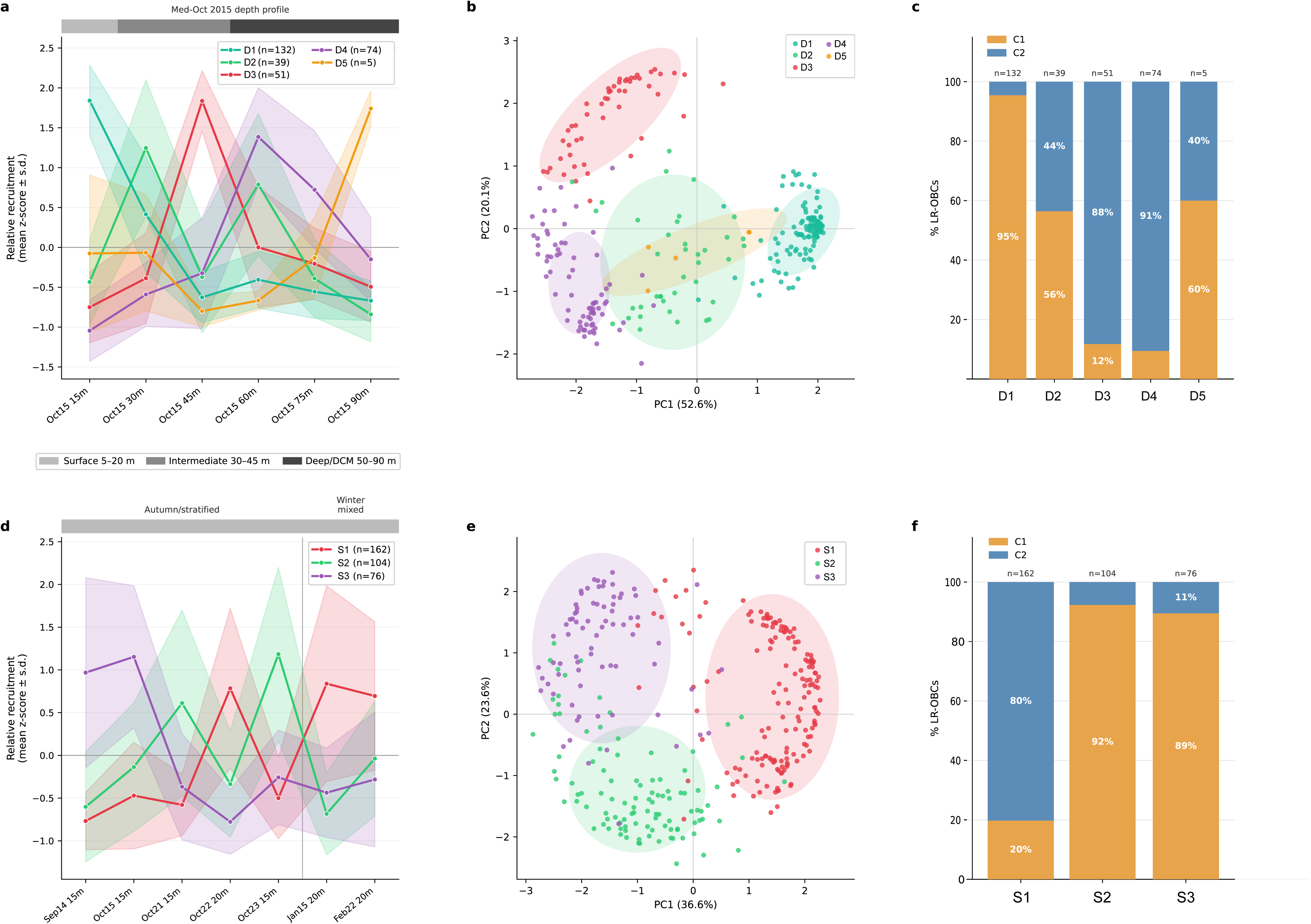
Hierarchical ecological structure of OBC-defined lineages within the gMED population. **a,** Mean relative recruitment profiles (mean Z-score ± s.d.) of five depth-specialized subpopulations identified by hierarchical clustering of the October 2015 vertical transect (301 LR-OBCs; k = 5, silhouette = 0.615). **b,** PCA of Z-score recruitment profiles for the October 2015 depth transect, coloured by subpopulation; shaded ellipses represent 1.8 standard deviations around each cluster centroid. **c,** Correspondence between depth-specialised subpopulations and global consortium assignment (C1, orange; C2, blue); numbers indicate sequence counts. **d,** Mean relative recruitment profiles of three temporal clusters identified from seven near-surface metagenomes (15–20 m) spanning 2014–2023 (342 LR-OBCs; k = 3, silhouette = 0.435). **e,** PCA of Z-score profiles for the near-surface time series, coloured by temporal cluster. **f,** Correspondence between temporal clusters and global consortium assignment, as in (c).

Both consortia were consistently recovered across the off-Alicante time series and the TARA Oceans Mediterranean stations, demonstrating that the covariation structure is not an artefact of a single sampling campaign or location. Importantly, sequence-sample origin was not included during clustering, yet origin categories showed a strong ecological association with consortium membership. LR-OBCs retrieved from DCM and lower-photic-zone metagenomes were enriched in C2, whereas OBCs originating from upper-photic and winter mixed-water collections were predominantly assigned to the C1 (Supplementary Fig.5 and Supplementary Table 4). Thus, an unsupervised analysis based only on recruitment covariation recovered the expected ecological stratification of the water column.

**Fig. 5:**
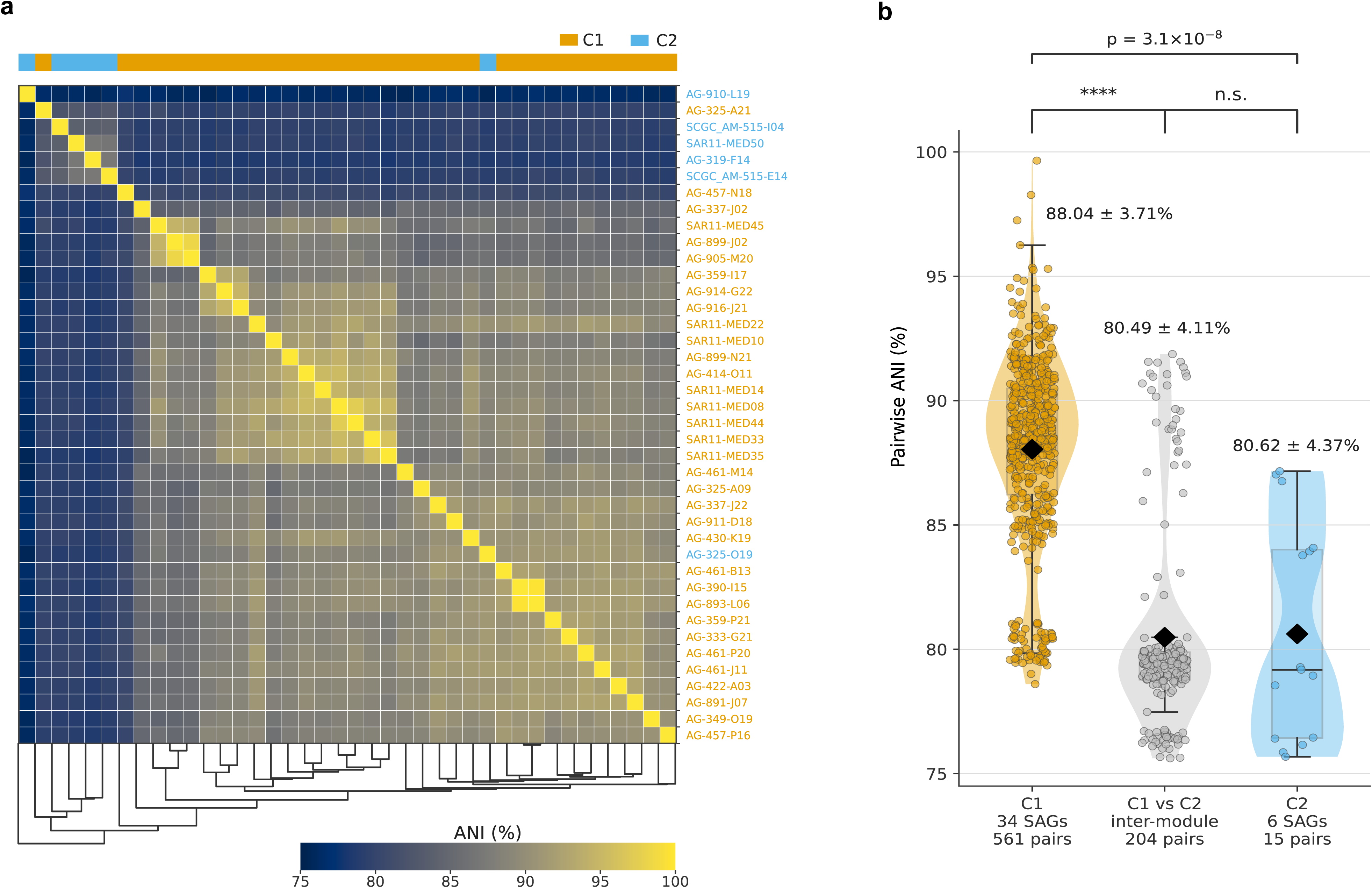
Core-genome ANI comparisons among gMED SAGs. **a,** Heatmap of pairwise average nucleotide identity (ANI) among 40 classified gMED SAGs ordered by hierarchical clustering. The colour bar at the top indicates consortium assignment (C1, orange; C2, blue). **b,** Violin plots of pairwise ANI distributions within consortium C1 (34 SAGs; 561 pairs), between consortia (C1 vs C2; 204 pairs), and within consortium C2 (6 SAGs; 15 pairs). Diamonds indicate median values; boxes show interquartile range. Statistical comparisons are indicated above (Mann–Whitney test).

The two-consortia partition also captures the dominant axis of ecological variation across the Mediterranean water column (Fig. 3a, lower panel). In stratified near-surface metagenomes (5–20 m), consortium C1 accounted for the overwhelming majority of total LR-OBC RPKG: 97.9% in Med-SEP2014 (15 m), 96.9% in Med-OCT2015 (15 m), 96.0% in Med-OCT2023 (15 m), 97.3% in TARA_009 (5 m), and 99.3% in TARA_030 (5 m). This near-complete dominance of the surface consortium under stratified conditions confirms that consortium C1 follows the upper-photic ecological niche with high fidelity. Three notable exceptions disrupted this pattern: the winter mixed-water samples (Med-JAN2015: C1 = 61.0%; Med-FEB2022: C1 = 60.8%; Med-OCT2022: C1 = 47.2%), where the deep/DCM consortium contributed substantially to surface recruitment. This is consistent with vertical mixing during winter and late-autumn, when thermocline breakdown homogenises lineages throughout the water column. The TARA_004 surface sample (Gulf of Cádiz, 5 m) also showed an anomalously elevated C2 contribution (27.6%), possibly reflecting Atlantic water mass influence or mixed hydrographic conditions at this Atlantic-Mediterranean transitional station. In deep and DCM metagenomes (40–60 m), C2 dominated the majority of samples: 72.1% in TARA_009 (55 m), 68.6% in Med-OCT2015 (45 m), 65.5% in Med-OCT2023 (60 m), 66.8% in TARA_004 (40 m), and 63.5% in Med-OCT2015 (60 m). Notable exceptions were three eastern Mediterranean TARA stations where C1 dominated even at depth (TARA_018 60 m: 85.7%; TARA_025 50 m: 62.3%; TARA_030 60 m: 94.9%), a pattern discussed further below.

To evaluate whether the ecological structure observed for LR-OBCs was also maintained when incorporating SAG-derived OBCs, we repeated the clustering analysis using the combined dataset (354 LR-OBCs and 31 SAG-OBCs) (Supplementary Fig. 6 and Supplementary Table 4). Following log(1+x) transformation, row-wise z-score normalization, correlation-based distances, and average-linkage hierarchical clustering, the optimal partition remained at two major consortia (silhouette = 0.289), whereas solutions with higher numbers of clusters showed substantially lower silhouette values (k=3, 0.258; k=4, 0.197; k=5, 0.170). The resulting clusters comprised 156 and 229 OBCs, respectively. Notably, SAG-derived OBCs did not form independent modules but were almost entirely embedded within one of the two major consortia, with 30 of 31 informative SAG-OBCs assigned to the surface-associated consortium C1 and only a single SAG-OBC assigned to the deep consortium C2. The consortium structure was highly robust to dataset composition as 352 of 354 LR-OBCs (99.4%) received identical assignments in the LR-only and joint analyses. The two OBCs that changed assignment — both of Winter type — displayed near-zero or negative silhouette scores in both analyses (silhouette = −0.017 and +0.135 in the LR-only analysis; −0.143 and +0.016 in the joint analysis), indicating that they occupy the boundary between consortia rather than representing genuine ecological reclassifications. These results indicate that the dominant ecological signal underlying OBC distribution is shared by both LR- and SAG-derived datasets and is largely captured by the same surface-versus-deep ecological gradient identified in the LR-OBC analysis.

Together, these results indicate that the gMED population of *Pelagibacterales* is not structured as a collection of hundreds of independently fluctuating OBC-defined lineages, but as two ecologically distinct and coordinated clonal consortia. Although depth-associated ecotypes have previously been recognised within *Pelagibacterales*, the structure described here differs in scale and meaning; it occurs within a single gMED genomospecies. It is resolved through O-chain biosynthesis clusters (i.e. variable flexomes), revealing coordinated modulation among closely related clonal lineages rather than divergence between phylogenetic units (see below). One consortium is associated with upper-photic conditions, whereas the other increases in relative importance in deep, DCM or vertically mixed waters. Thus, frequency modulation operates at the level of consortia rather than individual clones: the overall clonal repertoire remains highly diverse and persistent, while its internal composition is reorganised through coherent shifts in the relative abundance of surface- and deep/DCM-associated consortia across years, seasons, depths and geographical settings. This provides a population-level mechanistic model in *Pelagibacterales*, in which hydrographic change modulates the relative representation of alternative OBC-defined clonal consortia without losing the underlying population pangenome wealth. We term this model frequency modulation (FM).

### Depth and longitudinal structure of gMED consortia

To investigate whether such large-scale consortia may conceal additional structure associated with environmental gradients, we examined the internal organisation of the LR-OBC repertoire using both a high-resolution vertical transect from Alicante (October 2015) and geographically distributed Mediterranean metagenomes from the TARA Oceans expedition.

In the October 2015 depth profile (Haro-Moreno et al. 2018), 301 of the 354 LR-OBCs were recruited in at least two depth layers and were clustered according to their abundance-independent profiles (OBCs that recruited in only a single depth layer were excluded as their profiles lack the co-variation signal required for meaningful clustering). This analysis resolved five depth-specialised subpopulations (silhouette = 0.615; Supplementary Fig.4), each displaying a distinct depth optimum: 15 m (D1; n = 132, silhouette = 0.754), 30 m (D2; n = 39, silhouette = 0.303), 45 m (D3; n = 51, silhouette = 0.638), 60 m (D4; n = 74, silhouette = 0.520), and 90 m (D5; n = 5, silhouette = 0.536) (Fig. 4a,b; Supplementary Table 5b). The 15 m cluster was the most abundant and cohesive, accounting for 44% of all recruiting sequences. These depth-specialised subpopulations were non-randomly associated with the two major covariation consortia: LR-OBCs assigned to C2 were predominantly linked to clusters peaking at 45–60 m, whereas those assigned to C1 were concentrated in the 15 m cluster (Fig. 4c). The broad surface/deep partition therefore resolves, within a single water column sampled on a single occasion, into at least five distinct depth niches separated by intervals of just 15–30 m. This degree of fine-scale vertical partitioning among lineages within a single genomospecies suggests that OBC-defined clonal diversity encodes ecologically meaningful differences in depth preference, confirming depth as the primary driver of the population clonal composition.

To examine whether the two-consortium structure also manifests across temporal variation at a fixed depth, we applied the same clustering pipeline to the seven near-surface metagenomes (15–20 m) spanning 2014–2023 from the off-Alicante time series (342 LR-OBCs with recruitment in ≥2 samples; silhouette = 0.435; Supplementary Fig. 4). Three temporal clusters were identified (Fig. 4d–f; Supplementary Table 5c), S1 (n = 162, 80% C2-deep) peaked during winter and mixed-water conditions, consistent with the upward mixing of deep-associated lineages during thermocline breakdown; S2 (n = 104, 92% C1-surface) and S3 (n = 76, 89% C1-surface) both corresponded to stratified conditions but differed in their temporal peak, with S2 dominated by recent campaigns (2021–2023) and S3 by earlier ones (2014–2015). The same approach applied to the ten TARA Oceans Mediterranean metagenomes identified six geographically structured subgroups (silhouette = 0.382; Supplementary Figs. 4 and 7; Supplementary Table 5d), confirming that the surface/deep consortium signal extends across the broader Mediterranean basin with clusters dominated by surface samples being overwhelmingly C1 (98–100%) and the deep cluster 94% C2.

This east–west reversal in consortium dominance at depth might be a direct consequence of Mediterranean thermohaline circulation. Atlantic water entering through the Strait of Gibraltar flows eastward at the surface, progressively warming, evaporating, and gaining salinity. As it densifies, it deepens towards the Levantine basin, where it contributes to the formation of Levantine Intermediate Water (LIW). This progressive deepening of the water masses means that 60 m in the oligotrophic, ultra-stratified eastern Mediterranean corresponds ecologically to a much shallower depth in the western basin: the deep chlorophyll maximum (DCM) in the Levantine Sea lies at 80–120 m, compared to 40–60 m off Alicante (Lavigne et al. 2015). Consequently, a metagenome sampled at 60 m near Crete is still within the well-lit, nutrient-depleted upper photic zone that consortium C1 lineages exploit preferentially, whereas the same depth in the western basin falls clearly within the sub-euphotic conditions that favour consortium C2. The consortium boundary is therefore not defined by depth *per se*, but by the ecological conditions of light availability and nutrient supply — conditions that shift systematically eastward as Atlantic water descends along the Mediterranean thermohaline conveyor.

### The consortia do not reflect core-genome phylogeny

One possibility was that the consortia reflected a subspecies-level structure or another form of intraspecific taxonomy (e.g. ecotypes). If this were the case, genomes assigned to different consortia would be expected to show reduced average nucleotide identity (ANI) relative to genomes within the same consortium. We therefore used the available gMED SAGs containing sufficient core-genome sequence to calculate ANI values within and between consortia.

To assess genome-wide relatedness, we first reconstructed a whole-genome phylogeny of 48 gMED SAG genomes (Supplementary Fig. 8a). Clearly, C1 and C2 did not form reciprocally monophyletic groups but were interspersed throughout the tree, with C1 genomes distributed across multiple branches and the available C2 genomes failing to define an independent lineage. The ecological separation between surface-associated C1 and deep/DCM-associated C2 is therefore not explained by a reciprocal phylogenetic split in the conserved genome, indicating that consortium membership is superimposed on a shared core-genome background and is most likely encoded in flexible-genome content.

Nevertheless, pairwise ANI comparisons among classified gMED SAGs revealed that C1 SAGs were, on average, more similar to each other than to C2 SAGs: mean intra-C1 ANI was 88.04 ± 3.71% (34 SAGs; 561 pairs), compared with 80.49 ± 4.11% between consortia (204 pairs) and 80.62 ± 4.37% within C2 (6 SAGs; 15 pairs). Intra-C1 ANI was significantly higher than inter-consortium ANI (Mann–Whitney p = 3.1×10⁻⁸), whereas intra-C2 ANI was statistically indistinguishable from between-consortium values (p > 0.05), indicating that C2 does not constitute a genomically cohesive lineage distinct from C1. The broader ANI dispersion within C1 is consistent with its greater internal ecological sub-structure, as reflected in the five depth-specialised subpopulations identified within the surface consortium in the October 2015 vertical transect (Fig. 4a–c). Although the small number of available C2 SAGs precludes firm conclusions, the pattern is suggestive: the greater genomic diversity of C1 may reflect the more variable selective environment of the surface ocean, where intense fluctuations in irradiance, temperature and nutrient supply could favor a broader repertoire of genomic solutions, whereas the physically more stable deep photic zone only permits the coexistence of a genomically narrower lineage subset.

For the LR-OBCs, the available core-genome fraction was highly variable, making direct ANI comparisons difficult. However, the 16S–23S rRNA internal transcribed spacer (ITS), a marker widely used to resolve intraspecific diversity (Garcia-Martinez et al. 1999; Rocap et al. 2002), was available for a subset of LR-OBCs. We therefore reconstructed an ITS phylogeny including both LR-OBC- and SAG-derived sequences (266 sequences total), which allowed consortium assignments to be projected onto a marker-gene phylogenetic framework. Consistent with the SAG genome tree, C1 and C2 sequences did not form reciprocally monophyletic groups but were distributed throughout the tree without clustering by consortium membership (Supplementary Fig. 8b).

To quantify the degree of phylogenetic dispersion of consortium membership across both trees, we calculated the Association Index (AI) and Fitch Parsimony Score (PS) for the C1/C2 character, with significance assessed by permutation of consortium labels (n = 1,000). In the SAG genome tree, both statistics were significantly lower than expected under phylogenetic conservation of consortium membership (AI = 1.17, permuted mean = 3.99 ± 1.51, p = 0.017; PS = 3, permuted mean = 5.75 ± 0.48, p < 0.001). The same pattern was recovered in the ITS tree (AI = 12.25, permuted mean = 22.07 ± 2.32, p < 0.001; PS = 34, permuted mean = 48.28 ± 2.48, p < 0.001).

Together, these results confirm that consortium membership is not concordant with core-genome ancestry in either phylogenetic framework, indicating that the ecological differentiation between C1 and C2 is superimposed on a shared phylogenetic background and is most likely encoded in flexible-genome content and associated ecological functions, rather than in conserved genomic markers.

## Discussion

Building on the established diversity of *Pelagibacterales* gMED populations (Haro-Moreno et al. 2025; Molina-Pardines et al. 2026), we addressed the question of how stable this clonal diversity remains through time and across environmental gradients of depth and geography. The results indicate substantial persistence: clone abundances followed a log-normal distribution in all conditions examined, with a small number of lineages reaching high abundance, a larger number at intermediate levels, and a long tail of persistently low-abundance clones. This log-normal structure is the canonical signature of bet-hedging in community ecology, where it is interpreted as providing resilience through the maintenance of a reservoir of diverse, low-abundance genotypes. Transposed to the within-species level, the same logic applies: a genomospecies carrying high internal clonal diversity is better buffered against environmental change than one dominated by a single strain. The relative ranking of clones — their abundance hierarchy — was variable across samples, but rank shifts were frequently reversed rather than directional, and at no point did the data suggest clonal sweeps or succession over the time and environmental span considered. Taken together, these results support the proposal that clonal complexes act as coalitions that deploy their joint flexome repertoires to meet the challenge of dissolved organic carbon degradation (Rodriguez-Valera and Molina-Pardines 2026).

Unexpectedly, a deeper architecture was revealed beneath this stable diversity: the population harbours consortia — sets of clonal lineages that co-vary coordinately, as if the genomospecies possesses an internal community structure operating beyond the level of individual clones. At the Mediterranean basin scale, consortia C1 and C2 define a fundamental surface/deep ecological partition that is consistent across years and sampling locations, yet responsive to hydrographic state: the deep consortium invades surface waters during winter mixing, while the surface consortium penetrates deep layers in the oligotrophic eastern basin. This result demonstrates that the frequency modulation of gMED clonal consortia is directly coupled to basin-scale oceanographic forcing, providing a mechanistic link between Mediterranean thermohaline circulation and the relative representation of OBC-defined lineage subsets across the photic zone. At the local scale of a single water column, this partition further resolves into at least five depth-specialised lineage assemblages separated by intervals of only tens of metres — a degree of fine-scale ecological structuring that would be invisible to any analysis operating at the species or genomospecies level.

This hierarchical structure — global consortia, depth clusters, hydrographic modulation — provides a coherent ecological framework for the hundreds of OBC-defined clonal lineages maintained within the gMED population, and is consistent with a frequency modulation (FM) model in which environmental forcing controls the relative representation of lineage subsets rather than selecting for the wholesale replacement of one set of clones by another. Consortium modulation was most pronounced across depth gradients, with temporal and geographic variation having a lesser effect (depth > time > location). The FM analogy is deliberate: in electronics, frequency modulation encodes information in a carrier wave by varying its frequency rather than its amplitude; by analogy, the gMED population maintains a stable clonal pangenome — the amplitude — while the relative frequencies of individual lineage consortia fluctuate in response to environmental forcing. This frequency modulation reflects a necessary cooperation that keeps the system efficient and evolutionarily durable in the face of the chemical complexity of dissolved organic matter in aquatic environments (Dittmar 2021).

This work introduces a fundamental shift in our understanding of prokaryotic population genomics. The traditional view, derived from pure culture experiments in which a single clone multiplies in isolation, has long assumed the strain to be the functional unit of prokaryotes in nature — the microbial equivalent of the individual in animal or plant ecology. Here we show that, at least for the most abundant microbe in the ocean, this is not the case. The real functional and ecological units are clonal consortia comprising multiple lineages differentiated by their flexomes and the metabolic capacities these encode. The consortia comprise hundreds, if not thousands, of distinct clonal lineages whose combined pangenome exceeds the genome of any individual cell by orders of magnitude. Given the inherently small size of prokaryotic genomes, it is logical that only a combination of flexomes can provide the adaptive breadth required to meet the metabolic complexity that even the simplest natural habitat offers (Rodriguez-Valera and Molina-Pardines 2026). The flexome, not the core genome, is what determines the ecologically relevant differences among clonal lineages within a genomospecies.

One might argue that the consortia described here are simply a rebranding of the ecotypes discussed extensively in the literature (Cohan 2001). They are not. The critical distinction is phylogenetic: ecotypes, by definition, reflect core-genome divergence accumulated under differential selection, and are expected to form monophyletic or at least phylogenetically cohesive clusters. The consortia identified here show no such consistency — the core genome is relatively homogeneous across consortium members and does not covary with consortium assignment. This decoupling of ecological role from phylogenetic identity is precisely what the flexome-based consortium model predicts, and it is radically different from the classical ecotype framework.

The system analysed here — the photic zone of an oligotrophic sea — may not represent a universal microbial lifestyle. Recent culturomics studies of human microbiomes suggest that some body habitats harbour nearly clonal populations, with limited intraspecific diversity, albeit variable from host to host (Huang et al. 2023; Gichuki et al. 2025; Tripp et al. 2025; Pinedo-Bardales et al. 2026). However, animal-associated microbiomes differ fundamentally from free-living ones. Environments such as the gut or vagina are nutrient-rich and compositionally relatively simple — a point made plain by the ease with which their inhabitants grow in the laboratory, in stark contrast to the great plate count anomaly long documented in aquatic ecosystems. In such settings, a nearly clonal population suffices. For free-living microbes navigating the chemical complexity of marine dissolved organic matter, the picture is considerably more involved. We argue that the vast local pangenome of gMED reflects not merely an assembled genetic toolkit for the individual cell, but the collective functional repertoire of a population-level consortium — a public good maintained by phage predation, sustained by nutrient exchange through mechanisms such as extracellular vesicles, and shaped over evolutionary time by the irreducible complexity of the organic substrate (Rodriguez-Valera and Molina-Pardines 2026; Dittmar 2021).

These results reinforce the view that free-living prokaryotic genomes should be interpreted through the lens of the population rather than the individual cell. In nature, single-lineage populations are the exception, arising mainly in infectious diseases, specialised animal microbiomes, or industrial settings. For organisms such as gMED, ecological potential is not encoded in any single genome but is distributed across the collective pangenome of coexisting clonal lineages — a population-level architecture that provides the functional diversity, environmental resilience, and evolutionary stability revealed here. Complexity nested within complexity: a microbial analogue, perhaps, of the subatomic world revealed when physicists looked inside what had seemed to be elementary particles.

## METHODS

### Metagenomic datasets

Illumina short-read metagenomes from Tara Oceans, sampled in 2009 at stations TARA_004, TARA_009, TARA_018, TARA_025, and TARA_030 (Sunagawa et al. 2015), together with metagenomes from a fixed western Mediterranean station located 20 nautical miles off the coast of Alicante, Spain (37.354°N, 0.286°W), were used as recruitment databases (Haro-Moreno et al. 2018, Haro-Moreno et al. 2020, Haro-Moreno et al. 2025). The off-Alicante Mediterranean metagenomes span 2014–2023 and depths from 15 to 90 m, encompassing winter mixed-water and stratified summer/autumn conditions. Before recruitment, Illumina raw reads were quality-trimmed using Trimmomatic v0.39 with the following options: LEADING:3, TRAILING:3, SLIDINGWINDOW:4:15, MINLEN:50. Sample details and accession numbers are provided in Supplementary Table 1.

### OBC reference sequences

A total of 385 OBC sequences previously classified to belong to gMED were used as recruitment references, comprising 33 larger (five complete, “C”) SAG-derived OBCs (SAG-OBCs; mean length 24.4 Kb; range 6.0–67.7 Kb) and 354 partial OBC sequences recovered from PacBio CCS15 long reads (LR-OBCs; mean length 4.77 Kb; range 1.0–15.9 Kb) from five Mediterranean metagenomes (Haro-Moreno et al. 2025). Two SAG-OBCs form the original described collection, AG-430-E20 (1.1 Kb) and AG-914-O09 (2.2 Kb), were excluded owing to their anomalously short length to avoid artefactually inflated normalised prevalence values, reducing the initial set of 33 SAG-OBCs to 31.

Long reads were assigned to ecotype groups based on their sampling origin: Winter from mixed water-column samples; UP/UP2 from upper photic-zone samples during stratification; DCM from the deep chlorophyll maximum; and LP from the lower photic zone, following our previous classification. All OBC sequences were deposited under BioProject PRJNA1088973 in Haro-Moreno et al. (2025).

### Coverage calculation and normalisation

Illumina reads from each metagenome were recruited against the 385 OBC reference sequences using blastn v2.15 at 97% nucleotide identity and at least 90 bp of read alignment. Read recruitment was normalized by reference sequence length and metagenome size and expressed as RPKG values, calculated as recruited reads per kilobase of OBC sequence per gigabase of metagenomic data (Supplementary Table 2).

### Abundance-independent clustering of OBCs recruitment profiles

To identify groups of OBC-defined lineages that co-vary across metagenomes independently of their absolute abundance, we applied a row-wise Z-score normalisation to log-transformed RPKG values. Briefly, for each lineage *i* and metagenome *j*, the raw RPKG value *x*_ij_ was first transformed as *L*_ij_ = log(1 + *x*_ij_) to stabilise variance and handle zero values without pseudocounts. Each log-transformed row was then standardised as *z*_ij_ = (*L*_ij_ − *μ*_i_) / *σ*_i_, where *μ*_i_ and *σ*_i_ are the mean and standard deviation of lineage *i* across all metagenomes. This transformation preserves only the relative pattern of variation across metagenomes, making sequences with similar co-variation profiles comparable regardless of their absolute abundance.

Pairwise distances between lineage profiles were computed as correlation distances: *d*(*i*, *j*) = 1 − *r*(*z*_i_, *z*_j_), where *r* denotes the Pearson correlation coefficient. Hierarchical agglomerative clustering with average linkage was then applied to the resulting distance matrix. The optimal number of clusters *k* was selected by maximising the global silhouette coefficient *s*(*i*) = (*b*(*i*) − *a*(*i*)) / max(*a*(*i*), *b*(*i*)), where *a*(*i*) is the mean intra-cluster distance and *b*(*i*) the mean distance to the nearest neighbouring cluster, evaluated across *k* = 2 to 8. Biological interpretability and profile stability were also considered alongside the silhouette score. For each cluster, the mean Z-score profile and standard deviation across member lineages were computed and visualised. Principal component analysis (PCA) was applied to the Z-score matrix for a two-dimensional representation of global profile structure; clustering was performed exclusively in the full-dimensional space.

The global clustering analysis was applied to all 354 LR-OBCs across the complete set of 27 metagenomes, identifying two major covariation consortia (C1 and C2). To examine finer-scale ecological structure, the same pipeline was applied independently to two metagenome subsets. For the depth gradient analysis, only the six October 2015 metagenomes (15–90 m) were used; LR-OBCs were retained if they recruited in at least two depth layers (n = 301), as profiles with a single non-zero value lack the co-variation signal required for meaningful clustering. For the near-surface time series analysis, only the seven off-Alicante 15–20 m metagenomes spanning 2014–2023 were used; LR-OBCs were retained if they recruited in at least two of these metagenomes (n = 342). For the geographic analysis, only the ten TARA Oceans Mediterranean metagenomes (five surface and five deep/DCM samples from stations TARA_004, 009, 018, 025 and 030) were used; LR-OBCs were retained if they recruited in at least two of these metagenomes (n = 341). In all subset analyses, the Z-score normalisation, correlation distance, average-linkage clustering and silhouette-based k selection were applied identically to the global analysis, with k evaluated from 2 to 8.

All analyses were implemented in Python using NumPy, pandas, SciPy, and scikit-learn.

### Average nucleotide identity analysis

Pairwise average nucleotide identity, ANI, was calculated among all gMED SAG genomes using FastANI v1.3 (Jain et al. 2018) with default parameters. ANI comparisons were stratified into three groups: within-C1, within-C2, and between consortia (C1 vs C2). Differences between groups were assessed using the Mann–Whitney U test as implemented in SciPy v1.x. (Edwards et al.)

### Phylogenetic analysis and quantification of phylogenetic signal

To resolve the phylogenetic relationships among a large dataset containing 176 gMED SAGs and seven complete genomes of the Ia.1 genomospecies, a core-genome approach of shared proteins was employed using phylophlan v3.0 (Asnicar et al. 2020) with the following options: --database phylophlan –diversity medium –accurate. Muscle v3 (Edgar 2004) and trimAl (Capella-Gutierrez et al. 2009) were used for individual marker gene alignment and trimming of poorly aligned regions, respectively. The concatenated genome alignment was then used as the input for maximum-likelihood phylogenetic reconstruction using IQ-TREE v3 (Wong et al. 2026) with 5000 ultrafast bootstrap replicates and letting the program choose the best-fitted evolutionary model, which was VR+F+R5. For the ITS phylogeny, 273 gMED 16S–23S rRNA complete internal transcribed spacer sequences — comprising LR-OBC-derived and SAG-derived ITS *loci* were aligned using muscle and a neighbour-joining phylogenetic tree was constructed using MEGA 12 (Kumar et al. 2024) with 1000 bootstrap replicates, and with the Jukes-Cantor evolutionary model.

The degree of phylogenetic signal of consortium membership (C1 vs C2) was quantified using two complementary statistics: the Association Index (AI) and the Fitch Parsimony Score (PS). The AI was computed as the sum over all internal nodes of (1 − f_max)/(n − 1), where f_max is the frequency of the most common consortium state among the n classified leaves descending from that node. The PS was calculated using Fitch parsimony, counting the minimum number of C1↔C2 transitions required to explain the observed tip distribution. Statistical significance for both statistics was assessed by permuting consortium labels among classified sequences 1,000 times and computing the proportion of permuted values equal to or lower than the observed statistic (one-tailed test). All analyses were implemented in Python using NumPy and SciPy.

DATA: **Bioproject XXXXX.**

## Acknowledgements

This work was supported by grant “FLEX3GEN” PID2020-118052GB-I00 (co-funded with FEDER funds) from the Spanish Ministerio de Economía, Industria y Competitividad to F.R.-V., and by grant “SIMBAV” PID2024-162730NB-I00 from the Spanish Ministerio de Ciencia, Innovación y Universidades A.B.M.-C.). The authors declare no conflict of interest. The use of the large language model Claude (Anthropic) for some statistical analysis and language revision is acknowledged.

## SUPPLEMENTARY FIGURES

**Supplementary Fig. 1: Abundance distributions of OBC-defined clonal lineages across additional metagenomes. a,** Log₁₀-transformed RPKG abundance distributions of LR-derived OBCs across Mediterranean metagenomes not shown in Fig. 1. Histograms show the number of LR-derived OBCs within each abundance bin. Solid lines indicate fitted normal distributions in log₁₀ space and dashed vertical lines indicate the fitted mean. Statistics reported within each panel include the number of detected LR-derived OBCs, the mean (μ) and standard deviation (σ) of log₁₀-transformed RPKG values, the geometric mean abundance and the D’Agostino–Pearson normality test. Broad abundance distributions were consistently recovered across all metagenomes, indicating the coexistence of numerous OBC-defined clonal lineages spanning a wide abundance range. **b,** Log₁₀-transformed RPKG abundance distributions of SAG-derived OBCs across Mediterranean metagenomes not shown in Fig. 1. Histograms, fitted distributions and summary statistics are shown as in (a). Despite the smaller size of the SAG-derived reference set, abundance distributions closely resembled those obtained using LR-derived OBCs, supporting the robustness of the inferred clonal abundance structure. **c,** Log₁₀-transformed RPKG abundance distributions of LR-derived OBCs across TARA Ocean metagenomes. Histograms, fitted distributions and summary statistics are shown as in (a). Similar broad abundance distributions were recovered across geographically distant oceanic samples, indicating that the observed abundance structure is not restricted to Mediterranean populations. **d,** Log₁₀-transformed RPKG abundance distributions of SAG-derived OBCs across TARA Ocean metagenomes. Histograms, fitted distributions and summary statistics are shown as in (a).

**Supplementary Figure 2: Rank dynamics of SAG-derived OBCs across near-surface Mediterranean metagenomes. a,** RPKG-based rank trajectories of SAG-derived OBCs across seven near-surface Mediterranean metagenomes sampled between 2014 and 2023. Grey lines show all SAG-derived OBCs that reached the top-10 in at least one sample; coloured lines highlight representative stable and volatile lineages. The dashed line marks the top-10 threshold. **b**, Frequency of rank declines among SAG-derived OBCs entering the top-10 pool. Bars show the percentage of valid pairwise comparisons resulting in ≥3×, ≥5× or ≥10× rank declines. The inset shows the frequency of ≥3× rank declines stratified by starting rank. **c**, Relationship between recurrence and volatility for top-ranked SAG-derived OBCs. Recurrence is defined as the number of near-surface metagenomes in which each SAG-derived OBC entered the top-10, and volatility as the percentage of valid pairwise comparisons resulting in ≥3× rank declines. Grey points denote all other top-10 SAG-derived OBCs; coloured points indicate representative stable recurrent and volatile lineages.

**Supplementary Fig. 3: Two-consortium partition recovered from Mediterranean metagenomes excluding TARA Oceans stations. a,** Mean relative recruitment profiles (mean Z-score ± s.d.) of the two major LR-OBC consortia from hierarchical clustering of 17 off-Alicante Mediterranean metagenomes; depth categories are indicated above (light grey, surface 5–20 m; mid grey, intermediate 30–45 m; dark grey, deep/DCM 50–90 m). **b,** PCA of Z-score recruitment profiles coloured by cluster assignment (C1, orange, n = 198; C2, blue, n = 154); shaded ellipses represent 1.8 standard deviations around each centroid. **c,** Correspondence between cluster assignment in this restricted analysis and the global consortium assignment from the full 27-metagenome dataset.

**Supplementary Fig. 4: Silhouette coefficient as a function of cluster number across four independent sub-analyses.** Global silhouette coefficient for k = 2 to 8 clusters for four independent clustering analyses of LR-OBC recruitment profiles: depth gradient (October 2015 vertical transect, purple), near-surface time series 2014–2023 (orange), Mediterranean complete dataset without TARA (red), and TARA Oceans Mediterranean stations (teal). Optimal k values and corresponding silhouette scores are indicated for each analysis.

**Supplementary Fig. 5: Sequence origin and SAG-type composition of the two major consortia. a,** LR-OBC sequences assigned to both consortia (colours indicate sampling origin: upper photic zone stratified samples (UP, UP2), winter mixed-water samples (Winter), deep chlorophyll maximum (DCM), and lower photic zone (LP)). **b,** SAG-OBC clusters identified in the joint LR+SAG analysis (complete SAGs, C, dark blue; fragmented SAGs, F, light blue).

**Supplementary Fig. 6: Joint clustering of LR-OBC and SAG-OBC recruitment profiles. a,** Mean relative recruitment profiles (mean Z-score ± s.d.) of the two major consortia recovered from the combined dataset of 354 LR-OBCs and 31 SAG-OBCs across all 27 metagenomes. Depth categories: light grey, surface 5–20 m; mid grey, intermediate 30–45 m; dark grey, deep/DCM 50–90 m. **b,** Global silhouette coefficient for clustering solutions k = 2 to 10; the optimal partition is highlighted in orange. **c,** PCA of Z-score profiles for the joint dataset coloured by cluster assignment; circles denote LR-OBCs and triangles denote SAG-OBCs. **d,** Absolute counts of LR-OBC and SAG-OBC sequences assigned to each consortium.

**Supplementary Fig. 7. Geographic structure of gMED LR-OBC recruitment across TARA Oceans Mediterranean stations. a,** Mean relative recruitment profiles (mean Z-score ± s.d.) of six geographically structured subpopulations identified by hierarchical clustering of 341 LR-OBCs across ten TARA Oceans Mediterranean metagenomes from stations TARA_004, TARA_009, TARA_018, TARA_025 and TARA_030 (k = 6, silhouette = 0.382); samples are ordered from western to eastern Mediterranean with surface and deep/DCM layers shown separately. **b,** PCA of Z-score recruitment profiles coloured by geographic cluster; shaded ellipses represent 1.8 standard deviations around each centroid. c, Correspondence between geographic clusters and global consortium assignment (C1, orange; C2, blue); numbers indicate sequence counts per cluster.

**Supplementary Fig. 8. Phylogenetic trees of gMED genomes and ITS sequences. a,** Neighbour-joining phylogenetic tree of 273 gMED 16S–23S rRNA internal transcribed spacer (ITS) sequences, comprising 157 LR-OBC-derived and 107 SAG-derived sequences, aligned with MUSCLE v3 and constructed with MEGA 12 (1,000 bootstrap replicates; Jukes-Cantor model). **b,** Maximum-likelihood whole-genome phylogenetic tree of 176 gMED SAG genomes together with seven complete genomes of the Ia.1 genomospecies (outgroup), inferred using PhyloPhlAn v3.0 (--diversity medium --accurate) with MUSCLE v3 alignment, trimAl trimming, and IQ-TREE v3 reconstruction (5,000 ultrafast bootstrap replicates; best-fit model VR+F+R5). In both panels, branch and tip colours indicate consortium assignment based on OBC recruitment profiles: C1 (orange), C2 (blue), unassigned sequences (light grey), and reference/outgroup genomes (dark grey).

## SUPPLEMENTARY TABLES

**Supplementary Table 1.** Metadata of Illumina metagenomes used for OBC recruitment analysis.

**Supplementary Table 2.** RPKG-based recruitment matrix of OBC sequences across all metagenomes. **2A,** LR-OBC recruitment values. **2B,** SAG-OBC recruitment values.

**Supplementary Table 3.** Per-metagenome summary statistics of OBC recruitment. **3A,** LR-OBC recruitment summary. **3B,** SAG-OBC recruitment summary.

**Supplementary Table 4.** Consortium assignment, silhouette scores and Z-score recruitment profiles for all OBCs. **4A,** 354 LR-OBCs. **4B,** combined dataset of 354 LR-OBCs and 31 SAG-OBCs.

**Supplementary Table 5.** Cluster assignments for all 354 LR-OBCs across the global and subset clustering analyses. **5A,** Mediterranean-only analysis (k = 2). **5B,** October 2015 depth transect (k = 5). **5C,** Near-surface time series (k = 3). **5D,** TARA Oceans geographic analysis (k = 6).

## Notes

### Competing Interest Statement

The authors have declared no competing interest.

