## Supplementary figures for "Population dynamics of *Pelagibacterales* clonal lineages: ecological-consortia and frequency modulation"

**Supplementary Fig. 1a. Log(10) abundance distributions of LR-OCBs across Mediterranean metagenomes**

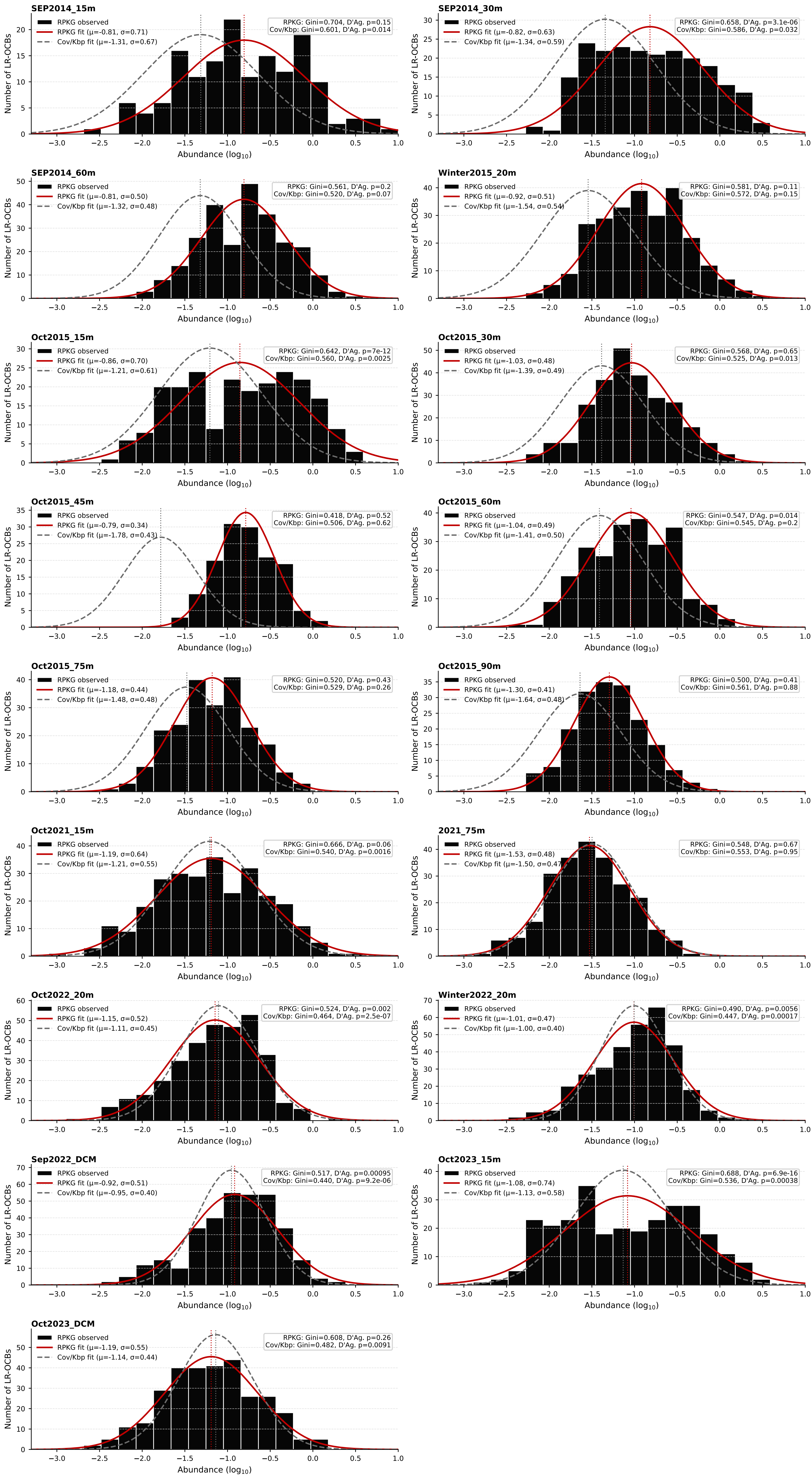

**Supplementary Fig. 1b. Log(10) distributions of LR-OCBs across TARA metagenomes**

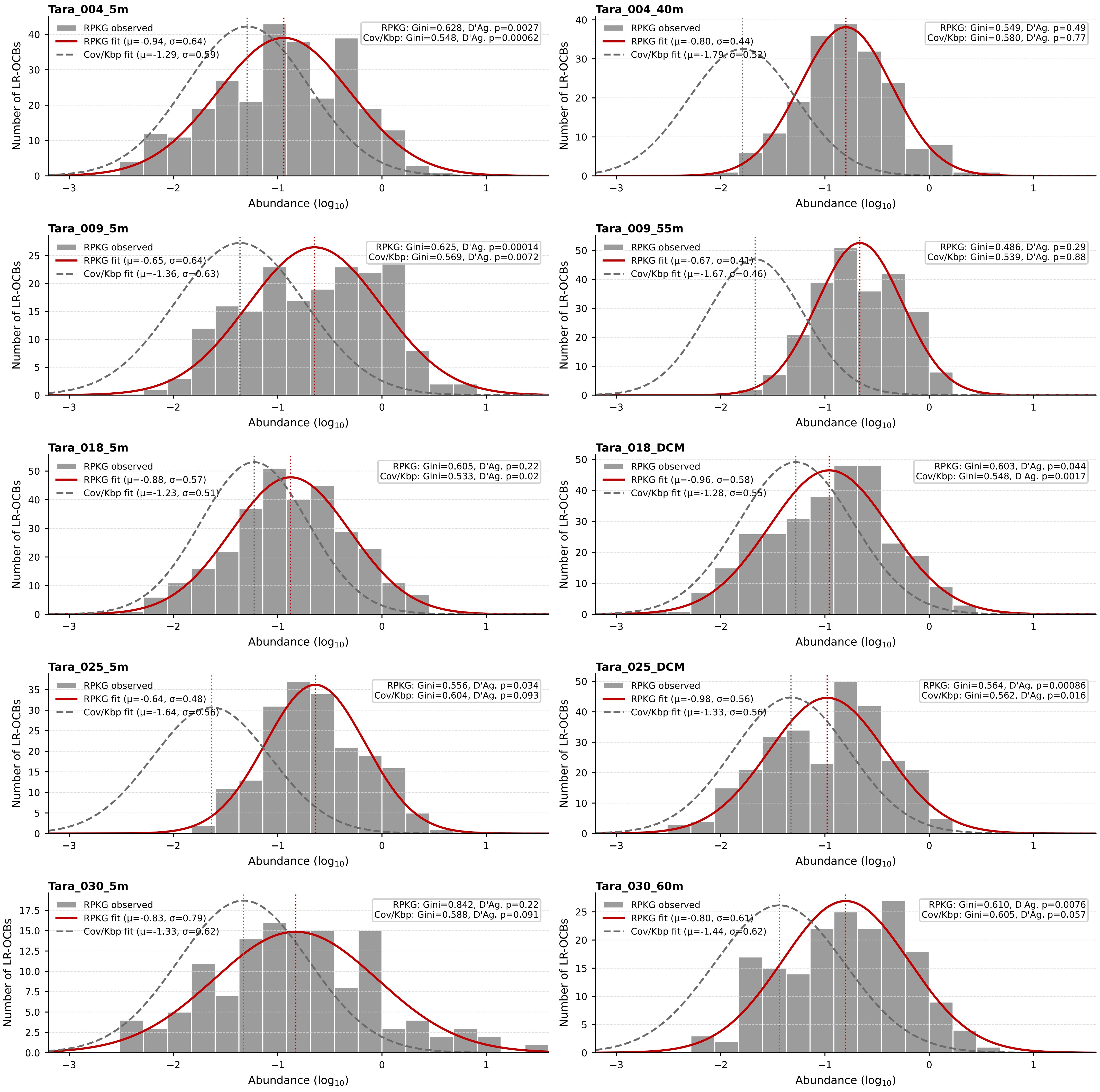

**Supplementary Fig. 1c. Log(10) abundancedistributions of SAG-OCBs across Mediterranean metagenomes**

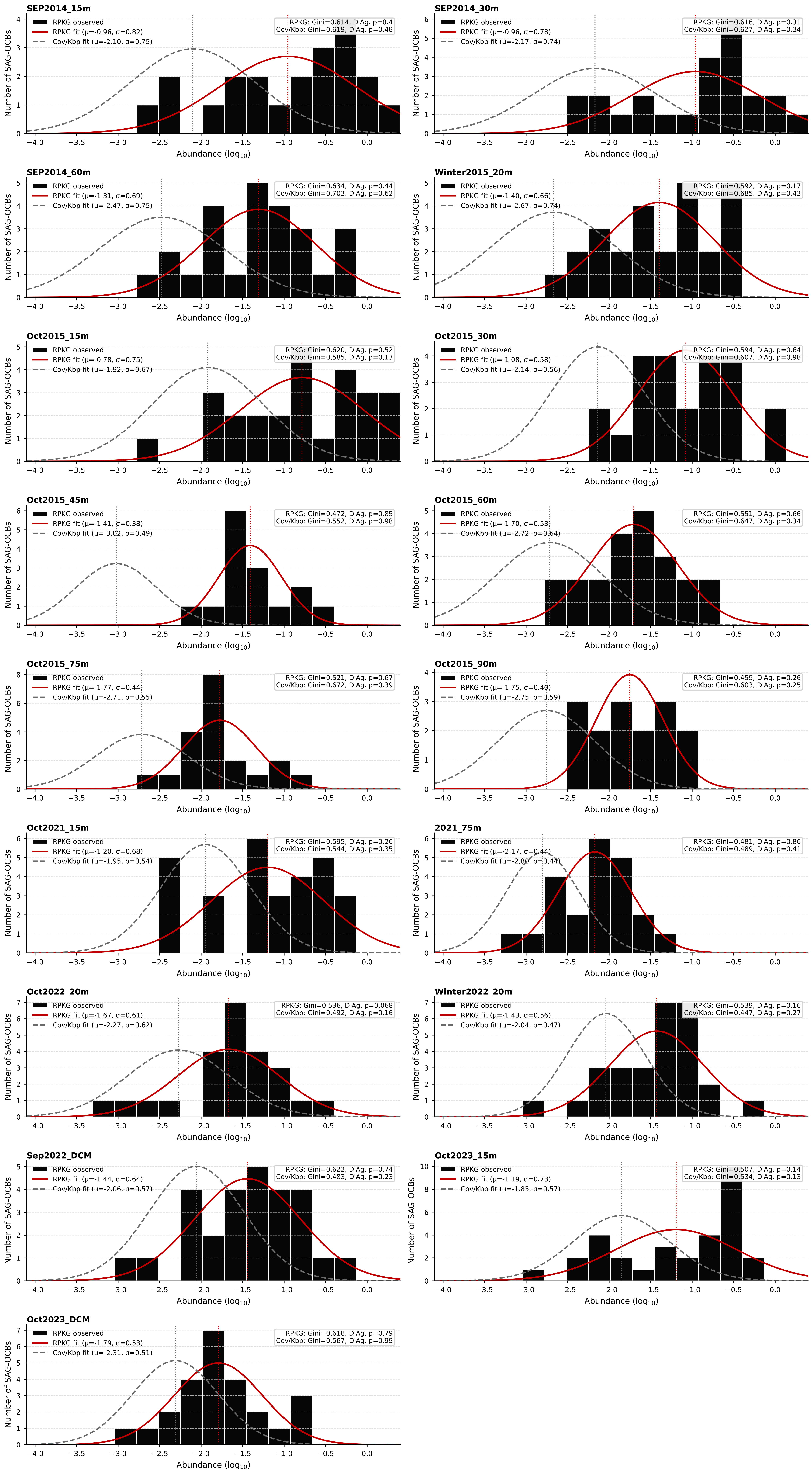

#### Supplementary Fig.1d. Log(10) abundance distributions of SAG-OCBs across TARA metagenomes

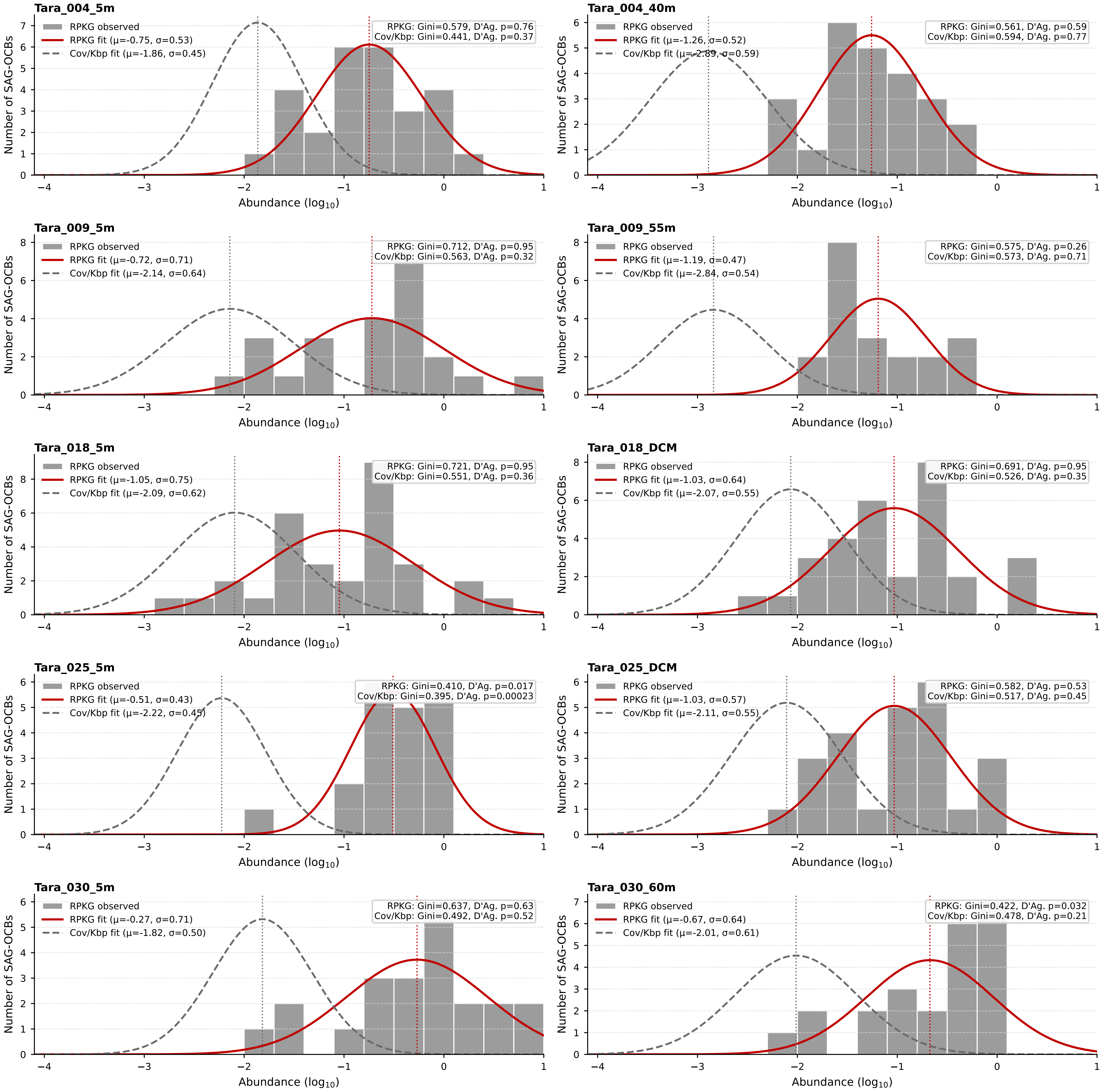

*Supplementary Figure 2*

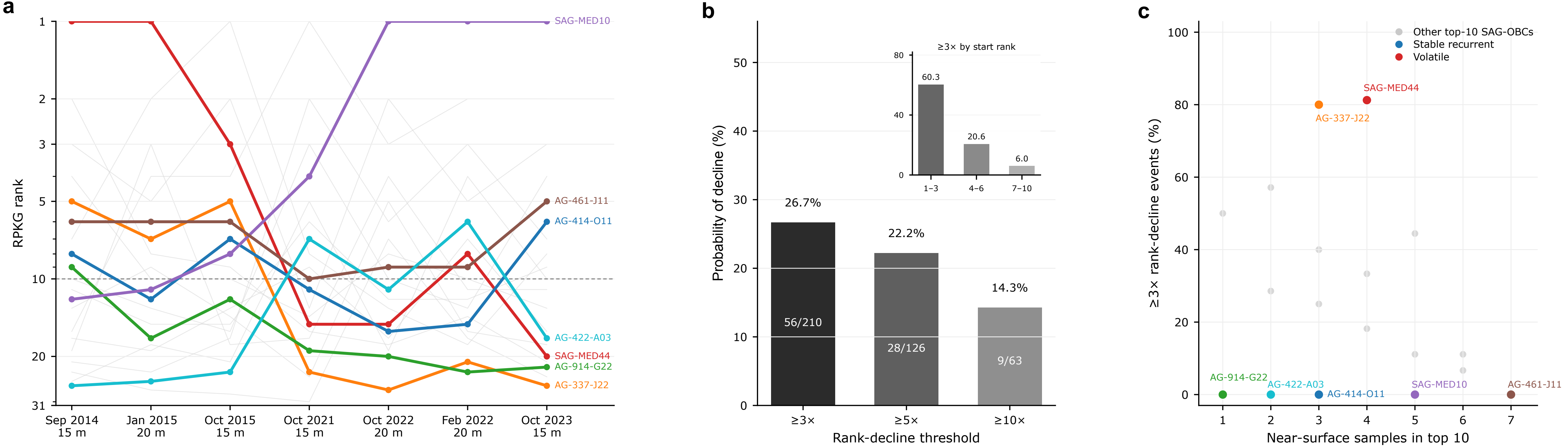

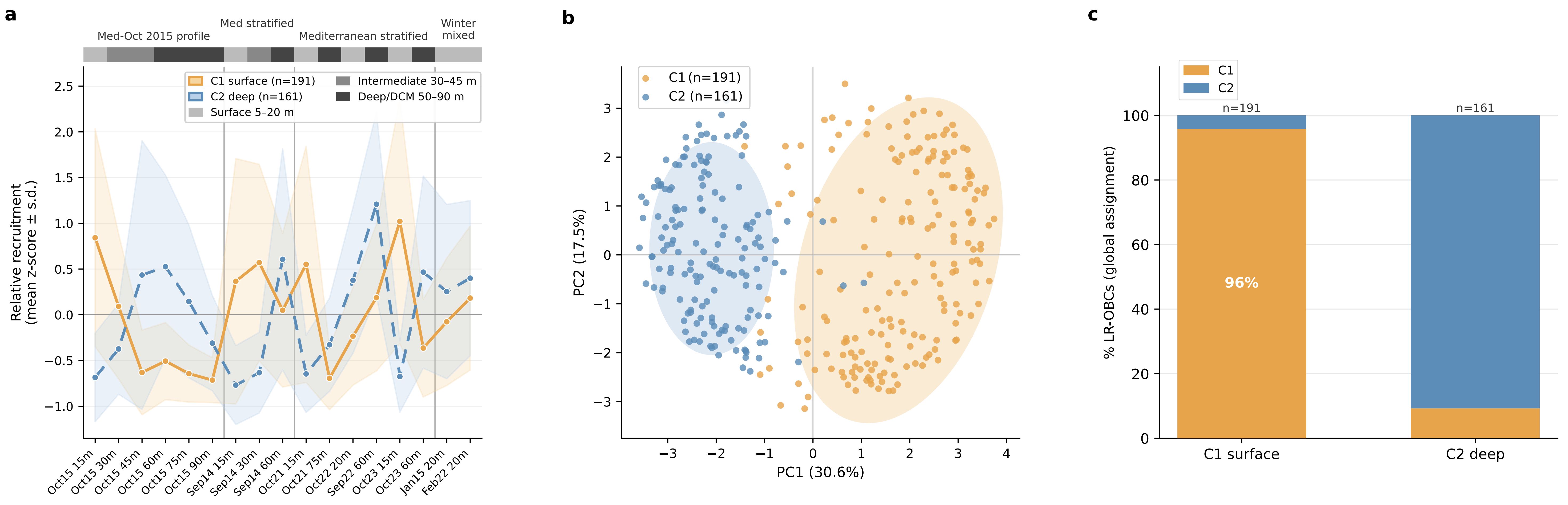


Supplementary Fig.3

Supplementary Figure 4

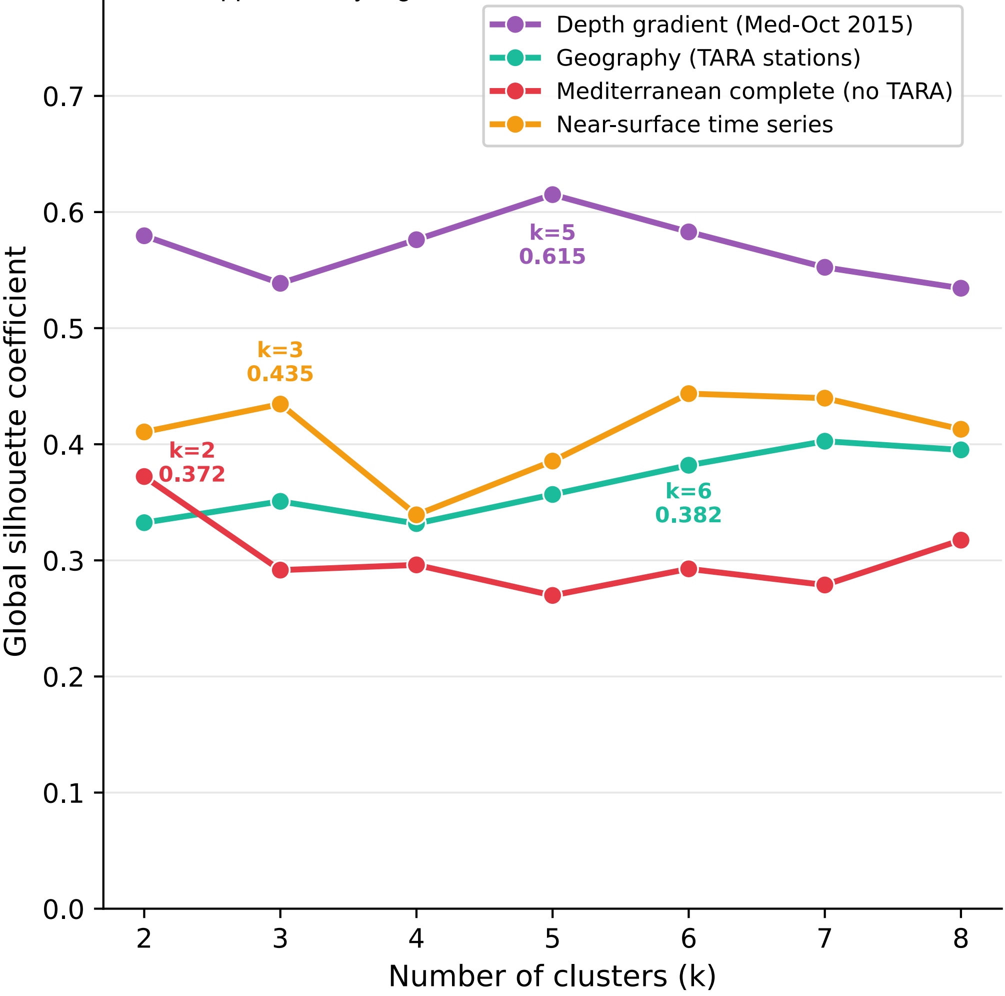

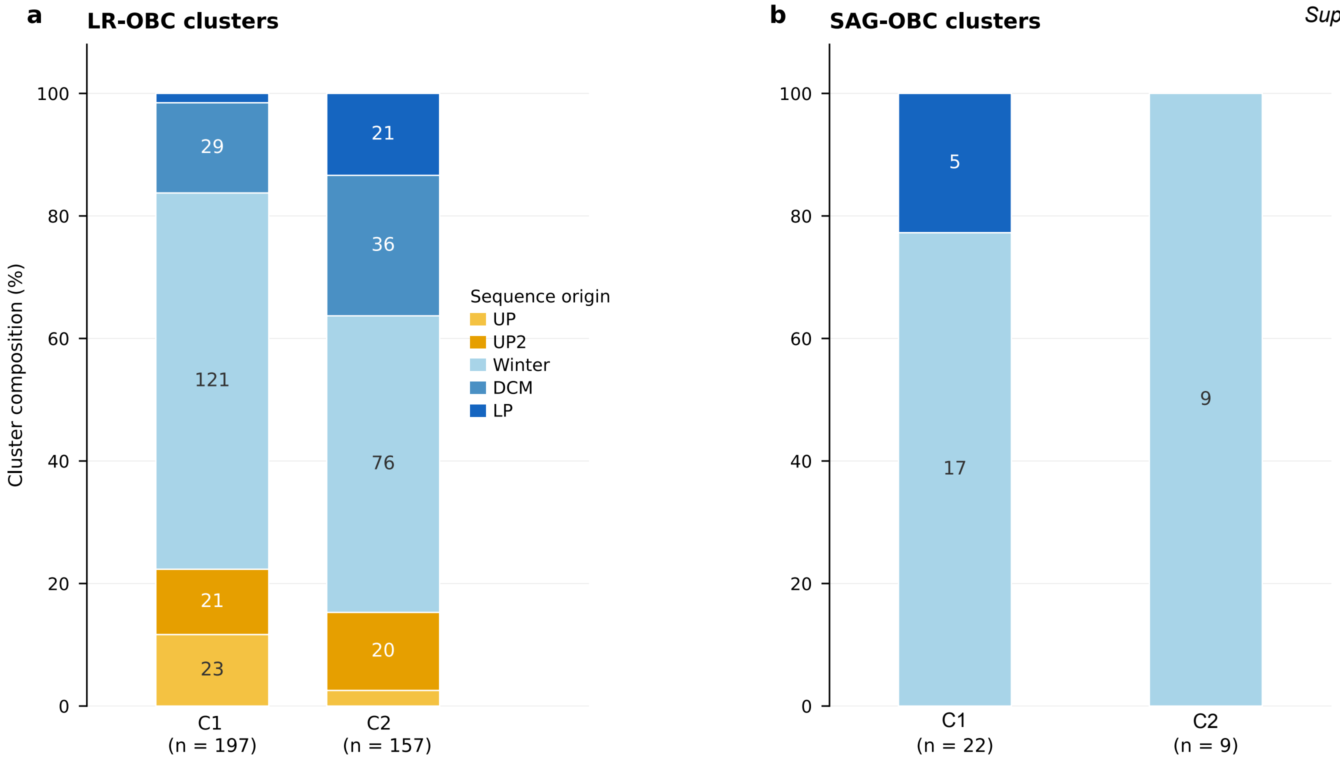
a
LR-OBC clusters
Cluster composition (%)
0
20
40
60
80
100
23
21
121
29
20
76
36
21
C1
(n = 197)
C2
(n = 157)
Sequence origin
UP
UP2
Winter
DCM
LP
b
SAG-OBC clusters
17
5
9
C1
(n = 22)
C2
(n = 9)

Supplementary Fig. 6

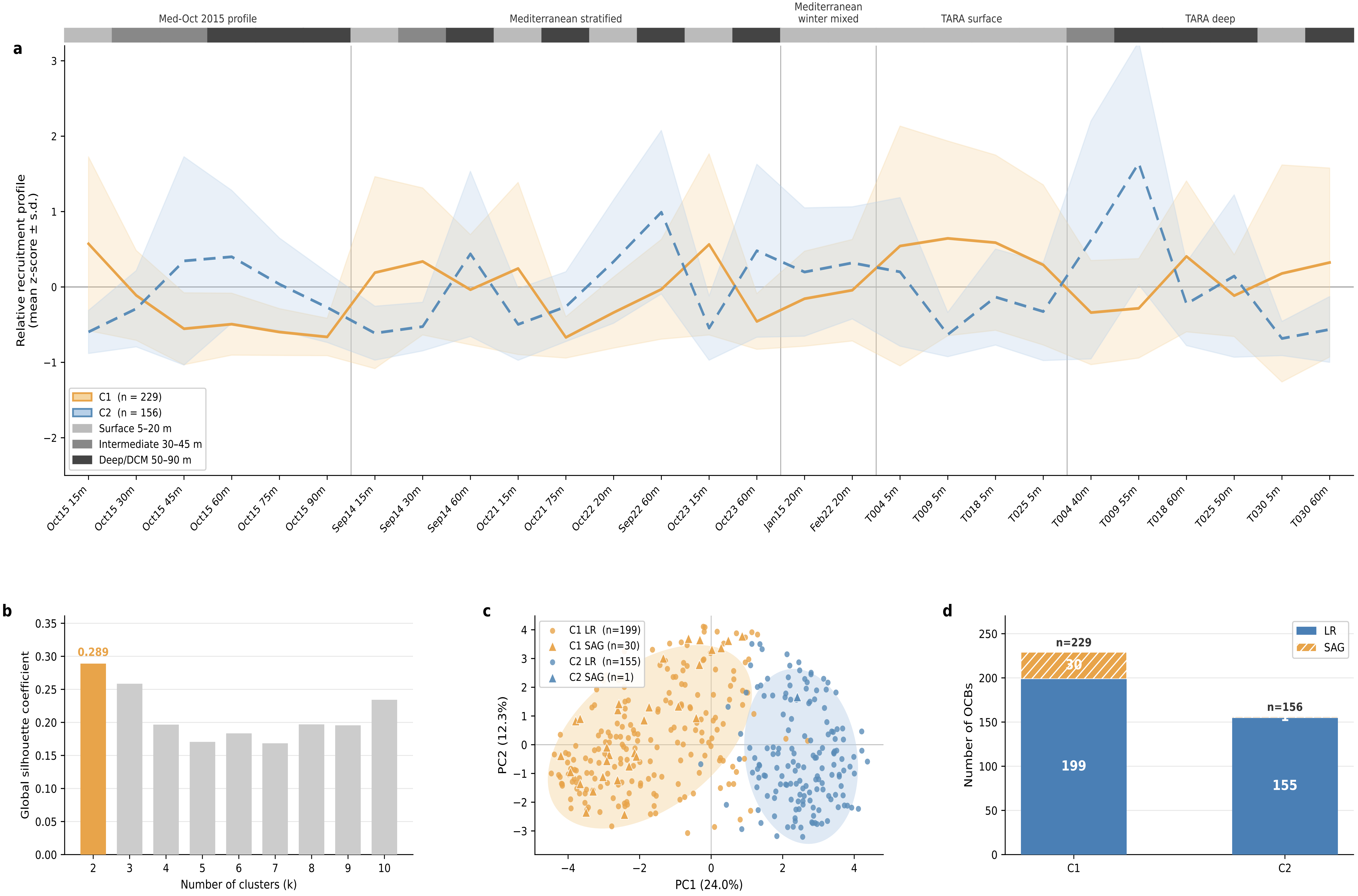

### Supplementary Fig.7

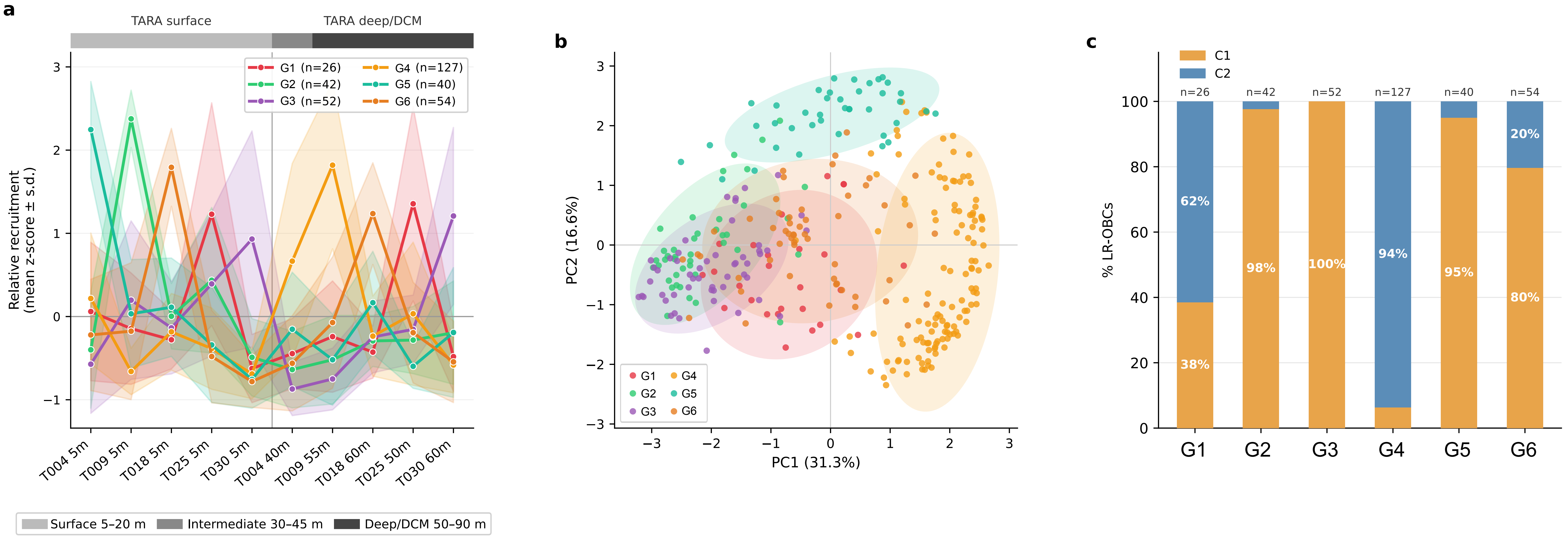

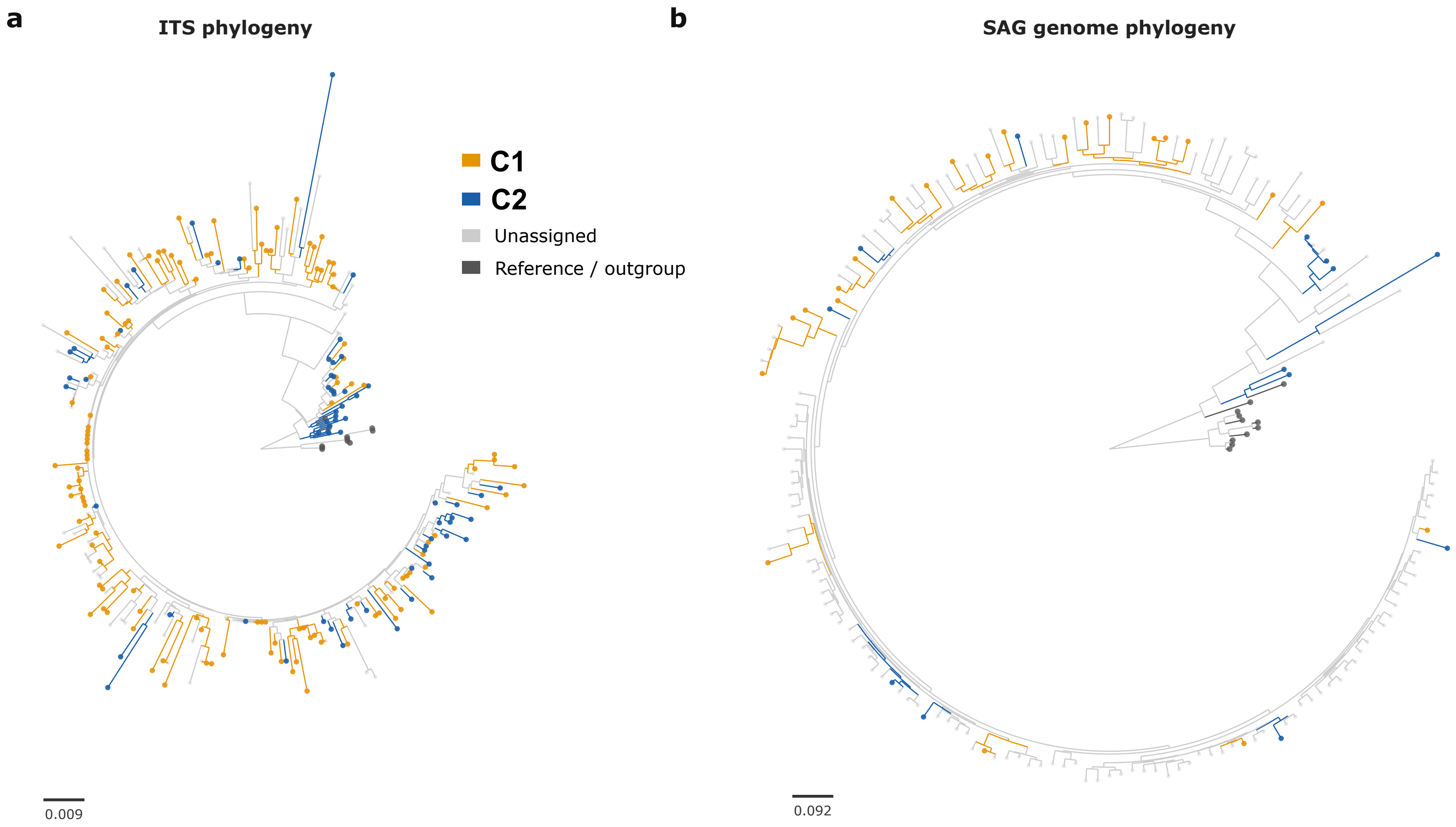
a
ITS phylogeny
b
SAG genome phylogeny
C1
C2
Unassigned
Reference / outgroup
0.009
0.092
